# The Loss of the E3 ubiquitin ligase TRIP12 inhibits Pancreatic Acinar Cell Plasticity and Tumor Cell Metastatic Capacity

**DOI:** 10.1101/2023.03.08.531649

**Authors:** Manon Brunet, Claire Vargas, Marjorie Fanjul, Laetitia Pieruccioni, Damien Varry, Guillaume Labrousse, Hubert Lulka, Florence Capilla, Anne Couvelard, Véronique Gigoux, Julie Guillermet-Guibert, Jérôme Torrisani, Marlène Dufresne

**Author notes:** <u>Correspondence:</u> Jérôme Torrisani, CRCT, 2 Avenue Hubert Curien, 31037 Toulouse. Marlène Dufresne CRCT, 2 Avenue Hubert Curien, 31037 Toulouse. contributed equally (Claire Vargas, Marjorie Fanjul). **<u>Grant Support:</u>** This project was funded by La Ligue Contre le Cancer, the Fondation Toulouse Cancer Santé (TCS2018CS079), l’Association Française pour la Recherche sur le Cancer du Pancréas (AFRCP) and Cancéropôle Grand Sud-Ouest (2021-EM22). Manon BRUNET was funded by the University Paul Sabatier and La Ligue Contre le Cancer. Claire VARGAS was funded by the University Paul Sabatier and La Ligue Contre le Cancer. Damien VARRY was funded by Ecole Normale Supérieure (Paris). The authors have nothing to disclose. RNA-Seq data: https://www.ncbi.nlm.nih.gov/geo/ accession number: GSE221587.

## Abstract

**Background & Aims:** Although specialized and dedicated to the production of digestive enzymes, pancreatic acinar cells harbor a high plasticity and are able to modify their identity. They undergo reversible acinar-to-ductal cell metaplasia (ADM) through epigenetic silencing of the acinar lineage gene program mainly controlled by PTF1a (Pancreas Transcription Factor 1a). ADM becomes irreversible in the presence of oncogenic Kras mutations and leads to the formation of preneoplastic lesions. We investigated the role of the E3 ubiquitin ligase Thyroid hormone Receptor Interacting Protein 12 (TRIP12), involved in PTF1a degradation, in pancreatic carcinogenesis.

**Methods:** We used genetically engineered mouse models of pancreas-selective Trip12 deletion, mutant Kras (G12D) and mutant Trp53 (R172H). We performed RNA sequencing analysis from acinar cells and cell lines derived from mice models tumors. We investigated the impact of TRIP12 deficiency on acute pancreatitis, tumor formation and metastasis development.

**Results:** TRIP12 is overexpressed in human pancreatic preneoplastic lesions and tumors. We show that a conditional deletion of TRIP12 in the pancreas during murine embryogenesis alters pancreas homeostasis and acinar cell genes expression patterns in adults. EGF induced-ADM is suppressed in TRIP12-depleted pancreatic acini. In vivo, a loss of TRIP12 prevents acini to develop ADM in response to pancreatic injury, the formation of Kras-induced pancreatic preneoplastic lesions, and impairs tumors and metastasis formation in the presence of mutated Trp53. TRIP12 is required for Claudin18.2 isoform expression in pancreatic tumors cells.

**Conclusions:** Our study identifies TRIP12 as a novel regulator of acinar fate in the adult pancreas with an important dual role in pancreatic carcinogenesis, in initiation steps and in metastatic behavior of tumor cells.

**Synopsis:** This study shows that Thyroid hormone Receptor Interacting Protein 12 plays an important dual role in the initiation steps and invasion of pancreatic carcinogenesis. Moreover, expression of TRIP12 switches on the expression of Claudin-18, a targetable biomarker of pancreatic tumors.

---

The mature exocrine pancreas comprises two types of epithelial cells. The acinar cells, the most abundant cell type, produce and secrete an extremely high level of digestive enzymes in pancreatic ducts daily. Acinar cells have been a model of terminal cellular differentiation with specialized cellular functions and organization dedicated to protein synthesis, transport, storage and exocytosis.^1^ As such, they were thought to be irreversibly differentiated. However, numerous studies revealed their surprising plasticity as a mechanism of defense against sustained stress as for example in the context of inflammation.^2–6^ Ductal cells were originally thought to be at the origin of pancreatic ductal adenocarcinoma (PDAC) due to ductal phenotype of pancreatic precancerous lesions and tumors. However, the nature of the cell at the origin of PDAC has been challenged and debated by experimental evidence providing insights in the plasticity of every pancreatic cell type (including endocrine cells) in the setting of neoplastic transformation. The most compelling studies used conditionally expressed mutant Kras^G12D^ allele combined or not with inactivation of tumor suppressor genes in one particular adult individual pancreatic cell type in mice.^7–12^ They demonstrated that cancer can originate from ductal and non-ductal cells of the mouse pancreas.

It has been known for decades that acinar cells lose their identity upon varying insults such as inflammation or in vitro culture through acinar-to-ductal metaplasia (ADM).^13–15^ During this process, acinar cells dedifferentiate to an intermediate state with embryonic features and may acquire mesenchymal properties.^3, 16^ Subsequently, acinar cells transdifferentiate to adopt a ductal phenotype. ADM is reversible, it can be induced by transient expression of reprogramming factors and it is coupled to the silencing of acinar cell-related genes.^17–20^ Numerous findings demonstrated that changes in acinar identity increase the susceptibility of cells to malignant transformation.^17, 21–24^ It is now widely accepted that acinar cells undoubtedly contribute to the development of pancreatic intraepithelial neoplasms (PanINs) preneoplastic lesions.

There is now a consensus on the importance of preserving a stable acinar cell differentiation program as a protective barrier against PDAC.^25–27^ Acinar cell differentiation program displays tumor suppressor properties by suppressing Kras function.^28^ Acinar cell-specific gene expression is driven by PTF1a (Pancreas Transcription Factor 1a) which is the main gatekeeper of acinar cell differentiation.^29, 30^ The silencing of acinar-cell related genes including *Ptf1a* is an initial epigenetic regulation step of ADM. ^17, 19, 20^ Maintaining or reintroducing PTF1a in the presence of Kras^G12D^ and inflammation can prevent or reverse tumor initiation.^28, 31, 32^

Thyroid hormone receptor interacting protein 12 (TRIP12) is an HECT (Homologous to the E6-AP Carboxyl Terminus) type E3 ubiquitin ligase. TRIP12 targets only a few substrates which exert critical functions in cell-cycle progression, DNA damage response, cell differentiation and chromatin remodeling.^33^ We previously demonstrated that TRIP12 is a chromatin associated protein regulating mitosis and that it targets PTF1a for proteasomal degradation.^34, 35^ We therefore hypothesized that TRIP12 plays a crucial role in PDAC and more particularly in regulating acinar cell fate in initiation steps of PDAC. We investigated the impact of conditional deletion of TRIP12 in mouse models for pancreatic regeneration, Kras^G12D^-mediated cancer initiation and progression.

## Results

### TRIP12 is overexpressed in human pancreatic cancers and preneoplastic lesions

We first investigated whether TRIP12 expression is associated with human pancreatic cancer. To this purpose, we performed an immunohistochemical analysis on human pancreatic tissue samples. TRIP12 is detected in the nucleus of exocrine acinar, ductal cells as well as endocrine islets of healthy pancreas. Nevertheless, TRIP12 immunostaining intensity remains weak and heterogeneous in healthy pancreatic tissue (Figure 1*A*). In contrast, we found that TRIP12 is markedly overexpressed in pancreatic adenocarcinoma (Figure 1*B*). This is supported by mRNA expression data from the Gene Expression Profiling Interactive Analysis (GEPIA) (Figure 1*C*). Interestingly, high Trip12 mRNA expression in primary tumors is associated with poorer patient survival, suggesting that TRIP12 plays a role during PDAC progression (Figure 1*D*).

**Figure 1:**
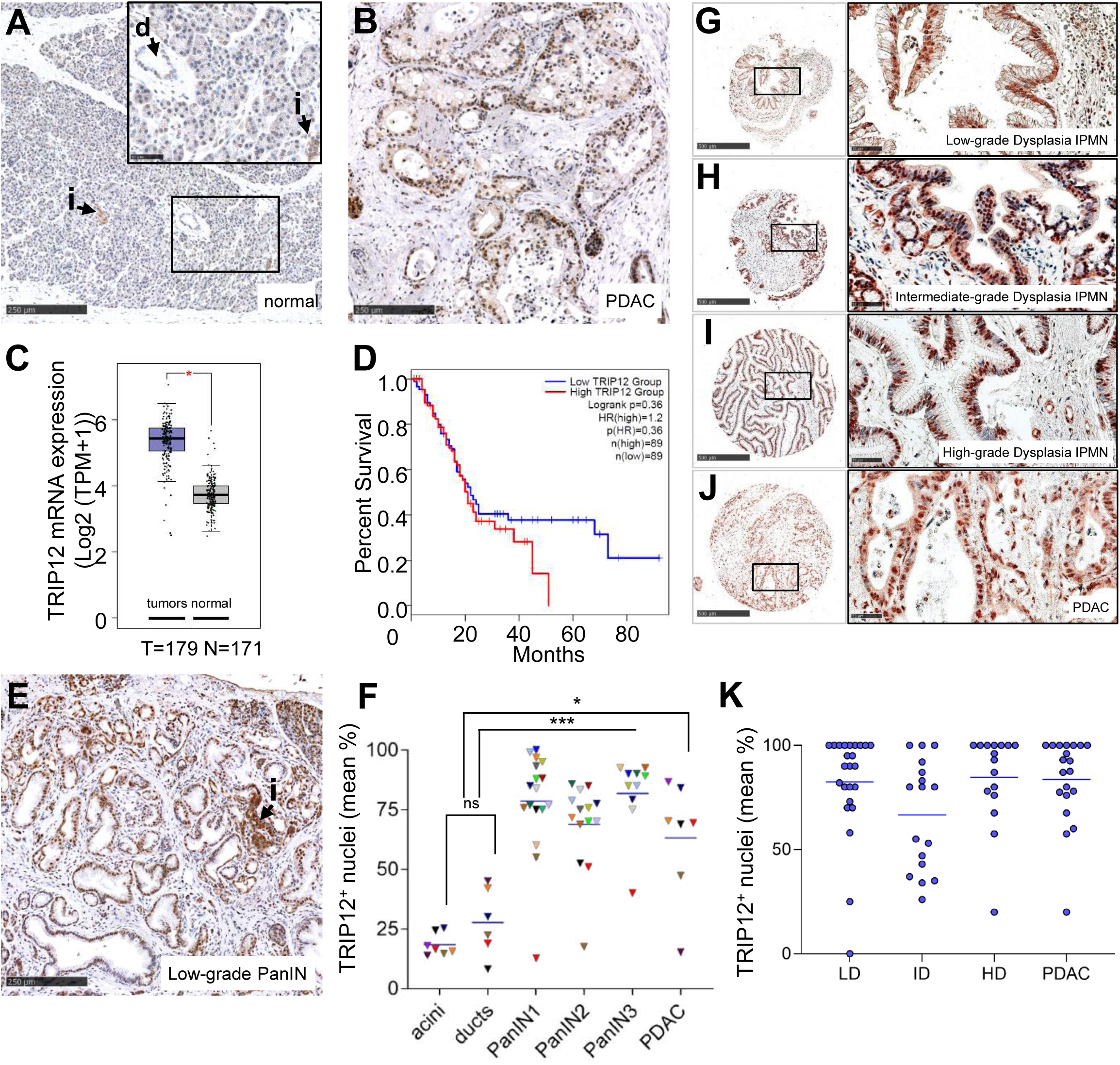
TRIP12 is overexpressed in human pancreatic cancers and preneoplastic PanIN and IPMN lesions. ***(A)*** TRIP12 immunohistochemistry staining on non-pathological pancreatic tissue (n=7), d=duct, i=islet. Scale bars: 250 μm (left), 50 µm (insert). ***(B)*** Representative TRIP12 immunohistochemistry staining in human pancreatic adenocarcinoma (n=7). Scale bar: 250 μm. ***(C)*** Expression of Trip12 mRNA in pancreatic tumors (purple, n=179) and normal pancreas (grey, n= 171), obtained from GEPIA database, *p<0.05 (unpaired t-test). ***(D)*** Kaplan Meyer survival plots of Trip12 mRNA high (red, n=89) and low (blue, n=89) expressing PDAC patients. Data were obtained from GEPIA database. **(*E*)** Representative TRIP12 immunohistochemistry staining in sections of human pancreas containing PanINs (n= 27). Scale bar: 250 μm. ***(F)*** Quantification of TRIP12 positive nuclei in 85 non neoplastic ducts, 29,000 non neoplastic acinar cells from 7 PDAC patients, in 170 PanIN1, 62 PanIN2 and 29 PanIN3 from 17, 14 and 10 patients, respectively and in 537 neoplastic ducts from 7 PDAC patients. Each symbol represents the mean percentage of TRIP12 positive nuclei in each quantified sample. Each color corresponds to one patient. The blue bars represent the grand mean of each quantified group. ***(G-J)*** Representative TRIP12 immunohistochemistry staining in IPMN: **G**, IPMN with low-grade dysplasia, **H**, IPMN with intermediate-grade dysplasia, **I**, IPMN with high-grade dysplasia, **J**, carcinoma. Scale bars of TMA cores are 500 µm and 50 µm for enlarged views. ***(K)*** Percentage of immunoreactive nuclei scored in 5 cores of 27 IPMN with low-grade dysplasia (LD), 17 IPMN with intermediate-grade dysplasia (ID), 20 IPMN with high-grade dysplasia (HD) and in 21 carcinoma (PDAC) by two investigators. The blue bars represent the grand mean of each quantified group.

Overexpression of TRIP12 is observed at the earliest stage of the disease in precancerous lesions such as in low grade PanINs (Figure 1*E* and *F*) and in intraductal papillary mucinous neoplasms (IPMN) with low-grade dysplasia (Figure 1*G-K*). This supports the hypothesis that TRIP12 plays an important role in the initiation steps of PDAC.

### TRIP12 deficiency during embryogenesis does not reveal major pancreatic abnormalities in embryos

To investigate the roles of TRIP12 in the initiation steps of PDAC, we decided to generate conditional Trip12 knockout mice. Since lethality was previously demonstrated in TRIP12 knock-in mutant mice at the embryonic stage E11.5, we investigated the functional requirement of TRIP12 during pancreas development.^36^ We first analyzed wild-type embryos at E13 and E15 and detected a low expression of TRIP12 in embryonic pancreas (Figure 2*A*). We then crossed mice carrying floxed alleles of Trip12 gene with Pdx1-Cre mice to deplete TRIP12 protein expression in embryonic pancreatic progenitors (Figure 2*B*). We examined whether the pancreas development is affected by the ablation of Trip12 gene in pancreatic progenitors using optical clearing of whole embryos for visualization of their pancreas at both embryonic stages. Quantification shows no difference in embryos and pancreas volumes between TRIP12-expressing controls C and TRIP12 knockout CT embryos (Figure 2*C*). The expression pattern of PDX1 staining in E13 TRIP12-deficient CT pancreas is similar to TRIP12-expressing controls (Figure 2*D*). We also found no difference in the expression pattern of PDX1, of PTF1a, and of amylase in the pancreas from E15 C and CT embryos (Figure 2*E*). This suggests that TRIP12 is dispensable for the setting of pancreas development in mice.

**Figure 2:**
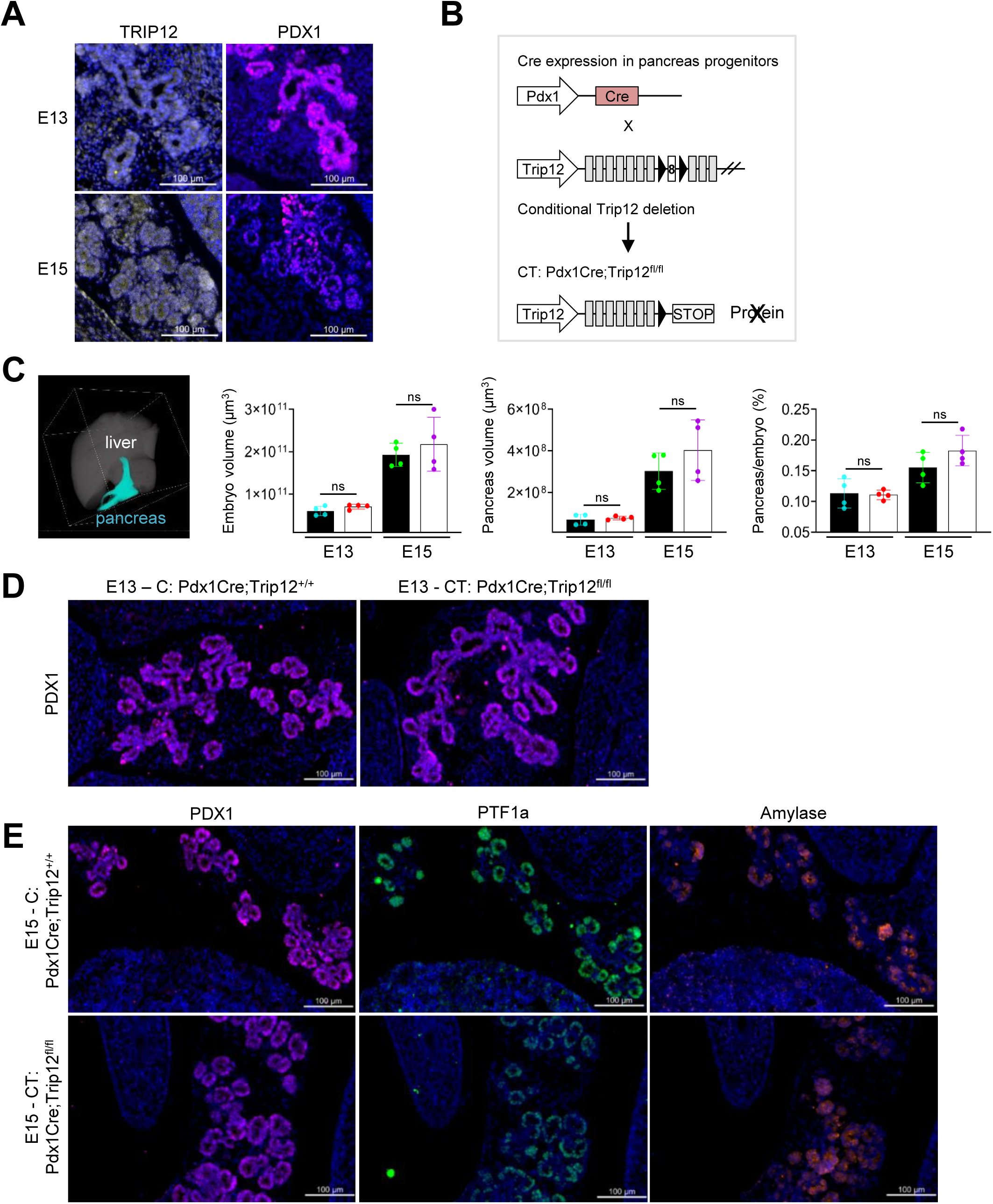
TRIP12 deficiency during embryogenesis reveals no major pancreatic abnormalities in embryos. ***(A)*** Immunofluorescence staining of TRIP12 (grey) and PDX1 (violet) in pancreas of E13 and E15 wild-type embryos. Nuclei are counterstained with DAPI. Scale bars: 100 μm. Representative images of three independent experiments are shown. ***(B)*** Schematic representation of the CT model in which the Cre recombinase expression is driven by the Pdx1 promoter. Black triangles indicate loxP sequences. ***(C)*** Quantification of the volume of C (black bars, n=4) and CT (white bars, n=4) E13 and E15 embryos and their pancreas. Embryos were optically cleared and acquisitions were performed with a selective plane illumination microscope (SPIM) technology. Segmentations were carried out as described in Methods using AMIRA software. Left panel:The segmented liver (grey) and pancreas (blue) of an C E13 embryo are represented in 3D by a volume rendering. The volume of the liver of each embryo was also determined and used as a control parameter unaffected by TRIP12 depletion. The bars represent the means ± SD. ***(D)*** Immunofluorescence staining of PDX1 in C and CT E13 embryos (representative of three independent experiments). Nuclei are counterstained with DAPI. Scale bars: 100 μm. ***(E)*** Immunofluorescence staining of PDX1 (violet), PTF1a (green) and amylase (orange) in C and CT E15 embryos (representative of three independent experiments). Nuclei are counterstained with DAPI. Scale bars: 100 μm.

### TRIP12 deficiency during embryogenesis reveals pancreatic abnormalities in adults

Pancreatic TRIP12-deficient CT mice are viable, appear healthy and are indistinguishable in appearance from the control littermates. We verified Cre-mediated recombination by crossing CT mice with ZsGreen Cre reporter mice to generate ZsG-CT mice (Figure 3*A*).^37^ ZsG-CT mice pancreas exhibit grossly normal morphology and as expected from PDX1-Cre-mediated recombination, ZsGreen is expressed in acinar cells (a), in ductal (d) and in islet cells. (Figure 3*A*).

**Figure 3:**
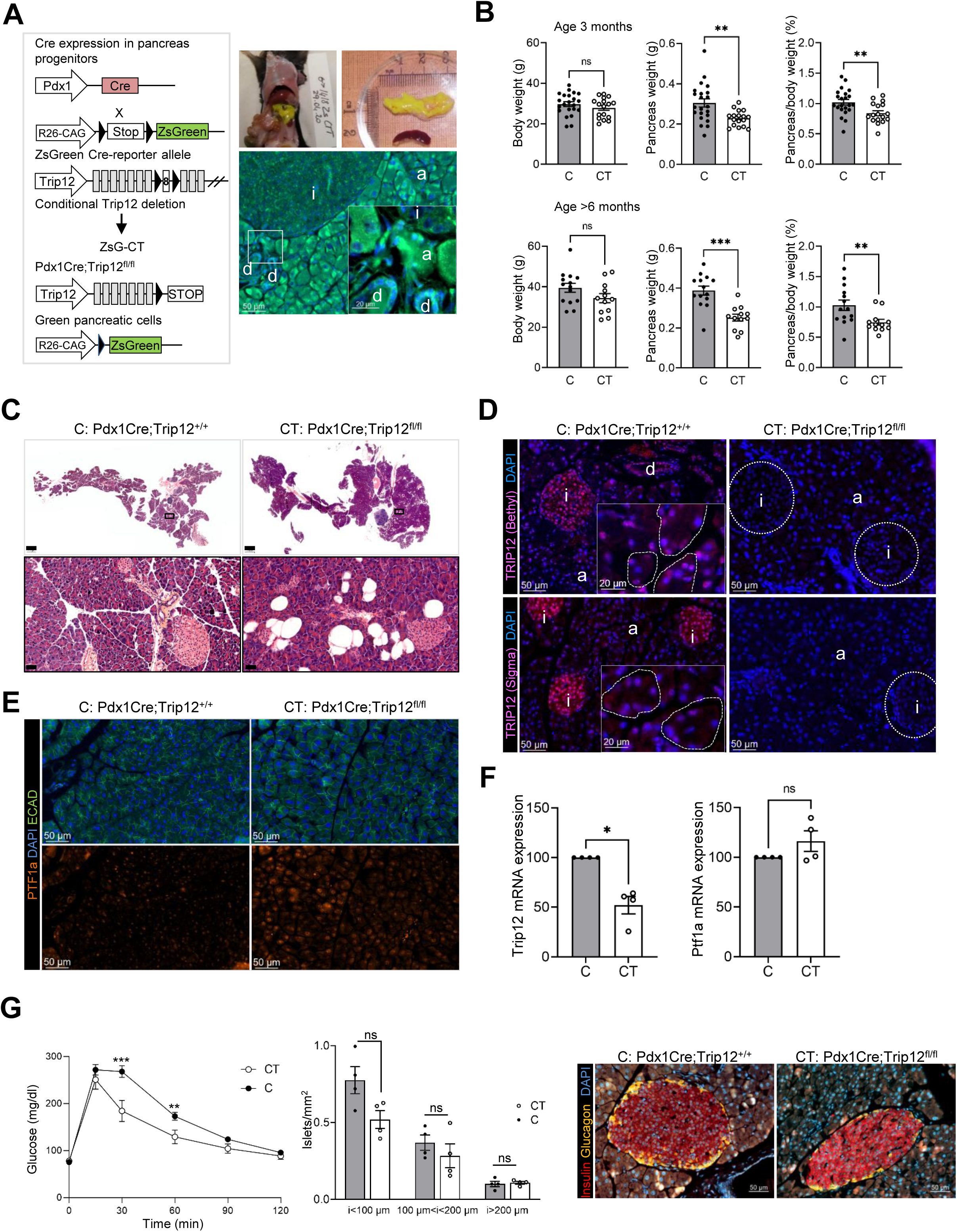
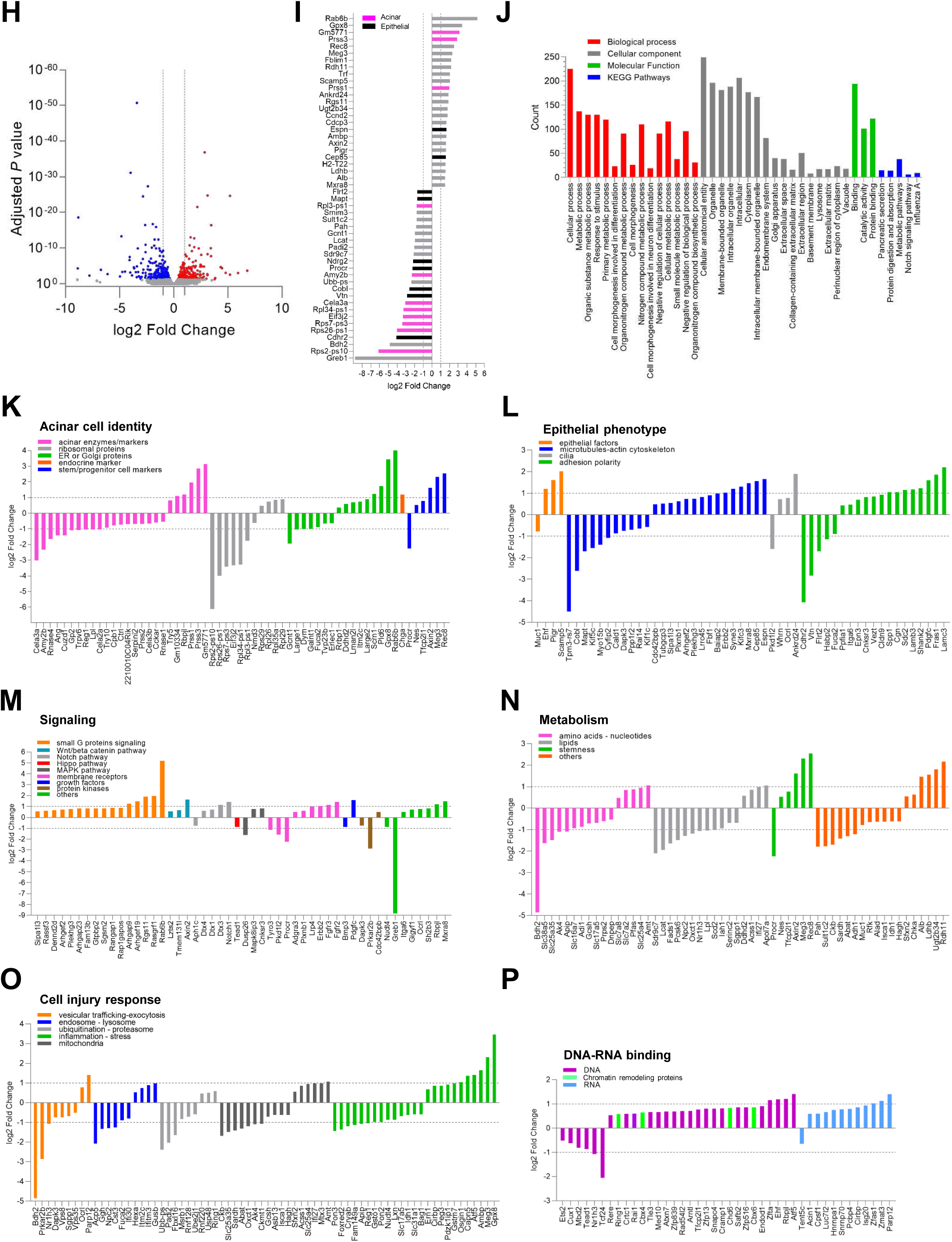
TRIP12 deficiency during embryogenesis reveals pancreatic abnormalities in adults. ***(A)*** Schematic representation of the Pdx1Cre;ZsGreen^f/f^;Trip12^f/f^ (ZsG-CT) mice and expression of ZsGreen in the pancreas of ZsG-CT mice showing recombination in all pancreatic cells; acinar cells (a), ductal cells (d) islet cells (i). Nuclei are counterstained with DAPI. Scale bars: 50μm, inset: 20 µm. ***(B)*** Body weight, pancreas weight and pancreas/body weight ratio of C control mice (n=22) and CT TRIP12 knockout mice (n=17) at the age of three months and of C (n= 12) and CT mice (n=14) older than six months of age. Data are expressed as mean ± SEM, **p<0.01, ***p<0.05, ns, non significant. ***(C)*** H&E staining of Pdx1Cre;Trip12^+/+^ (C) and Pdx1Cre;Trip12^fl/fl^ (CT) pancreas at the age of three months, scale bars: 1000 µm. Boxes indicate enlarged regions, scale bars: 50 μm. ***(D)*** Immunofluorescence staining of TRIP12 in C mice pancreas in acinar (a), ductal (d) and endocrine islet cells (i) using Bethyl anti-TRIP12 antibody (upper panels) and Sigma anti-TRIP12 antibody (lower panels). Images are representative of three independent experiments. Nuclei are counterstained with DAPI. Scale bars: 50 μm. insets, scale bars: 20 µm; a: acini, d: duct, i: islet. ***(E)*** Immunofluorescence staining of PTF1a (orange) in C and CT pancreas mice at the age of three months. Nuclei are counterstained with DAPI (blue). E-Cadherin staining (green) delimits the cells. Images are representative of three independent experiments. Scale bars: 50 μm ***(F)*** qRT-PCR analysis of TRIP12 and PTF1a mRNA levels in C (grey bars) and CT (white bars) isolated acinar cells (n=3). Data are expressed as a percentage of expression in control acinar cells normalized by HPRT, cyclophilin A and 18S genes expression, *p<0.05. ***(G)*** Left panel: glycemia after intraperitoneal glucose injection in control C (black circles, n=21) and CT TRIP12 knockout (white circles, n=15), mice were one to three months old, ***p<0.001, **p<0.005. Islets were counted and their diameter measured in sections of pancreas of C (grey bars and black circles, n=4), 2 sections per pancreas) and CT (white bars and white circles, n=4), 2 sections per pancreas. Data are expressed as mean ± SEM, ns, non significant. Right panel: representative immunofluorescence staining of insulin (red) and glucagon (yellow) in islets from C mice and CT mice (n=3). Nuclei are counterstained with DAPI. Scale bars: 50 μm. ***(H)*** Volcano plot of genes with differential expression in CT acinar cells from RNA-Seq of three biological replicates for C and CT pancreatic acini. Blue dots represent genes with a reduced expression, red dots represent genes with an increased expression, grey dots represent genes with an adjusted P value lower than 0.05. ***(I)*** Plot of the 25 most significantly increased or decreased mRNA in CT acinar cells; genes with an adjusted P value lower than 0.05 were considered statistically significantly altered, pink bars: genes related to acinar cell identity, black bars: genes related to epithelial phenotype. ***(J)*** Bar graphs representing a gene ontology analysis of biological process, cellular component, molecular function and KEGG pathways. ***(K-P)*** Changes in the expression of genes for the acinar cell identity (*K*), epithelial phenotype *(L*), signaling (*M*), metabolism (*N*), cell injury response (*O*), and DNA/RNA binding proteins (*P*).

While body weight of CT and control mice are similar at the age of three months, we observed significant alterations of pancreas weight and pancreas to body weight ratio in CT mice that persist in older mice (Figure 3*B*). The comparison of H&E staining shows a pancreatic steatosis in all TRIP12-deficient CT mice pancreas examined at the age of 3-4 months but no other cytological and degenerative changes (Figure 3*C*). We examined the pattern of TRIP12-expression in the murine pancreas in C and CT mice using two different antibodies raised against either a N-terminal peptide or a peptide located in the C-terminal part of TRIP12. Immunofluorescence staining confirms that similarly to human pancreas, TRIP12 is heterogeneously present in the nucleus of murine acinar, duct and islet cells with a subset of cells that do not express TRIP12 (Figure 3*D*). The loss of TRIP12 is visualized in all pancreatic cell types of CT mice pancreas (Figure 3*D*). Absence of staining using TRIP12 antibody recognizing the N-terminal sequence of the protein shows that TRIP12 N-terminal part of the protein is not expressed in CT. In parallel, immunofluorescence analysis reveals an increased PTF1a expression in TRIP12-deficient CT acinar cells compared to control C acini (Figure 3*E*). This is in accordance with our previous results that demonstrate a PTF1a targeting by TRIP12 and therefore validates the pancreatic knockout of Trip12.^34^ In support of a post-translational action of TRIP12 on PTF1a levels, PTF1a mRNA expression in acini is unaffected by the ablation of TRIP12 (Figure 3*F*).

We performed intraperitoneal glucose tolerance tests to evaluate whether conditional TRIP12 depletion in islets affects blood glucose homeostasis. Fasting glycemia is not significantly different in TRIP12-deficient CT compared to control mice at the age of three months (Figure 3*G*). However, the response to glucose overload is improved indicating that TRIP12 gene depletion in the pancreas affects pancreatic endocrine function of young adults with no significant difference of islets size, density, insulin and glucagon staining (Figure 3*G*).

We further studied the impact of TRIP12 knockdown by a RNA-Seq analysis of RNA extracted from TRIP12-deficient and control acinar cells. A total of 566 genes is significantly altered (adjusted P value ≤ 0.05) (Figure 3*H*). The top 25 most down regulated genes in CT acinar cells supports a role of TRIP12 in acinar cell phenotype maintenance since eight of these genes are related to the production of enzymes (amylase, elastase, five ribosomal proteins and eukaryotic translation initiation factor 3, subunit J2) and seven to the regulation of differentiation (cadherin related family member 2, vitronectin, Cordon-Bleu WH2 Repeat Protein, protein C receptor, N-Myc downstream-regulated gene 2 protein, microtubule-associated protein tau, fibronectin leucine rich transmembrane protein) (Figure 3*I*). Analysis of biological processes, cellular component and molecular function categories revealed changes in genes associated with metabolic and cell morphogenesis, organelles, vacuoles, extracellular matrix, pancreatic secretion and Notch pathway (Figure 3*J*). We therefore identified an effect of TRIP12 depletion on genes related to acinar cell identity, epithelial phenotype, signaling, metabolism, cell injury response, and nucleic acids binding proteins (Figure 3*K-P*).

Strikingly, nineteen secretory enzymes and several ribosomal proteins have a reduced gene expression. Not all acinar secretory enzymes genes are decreased since the expression of four genes of the trypsin family is increased together with the one of the pancreatic acinar specific transcription factor RBPJ-L (Figure 3*K*). Expression of the endocrine cells marker chromogranin A is also upregulated as well as the one of stem cells or progenitor markers such as nestin further indicating an altered acinar cell differentiation. In line with a modified acinar cell identity, several genes related to epithelial character are modulated (Figure 3*L*). The upregulation of several actors of small G proteins signaling genes supports the idea that many cellular functions are modified in TRIP12-deficient acinar cells (Figure 3*M*). Of note, four members of the Notch signaling cascade are also upregulated. Expression of mediators of amino-acids, nucleotides and lipids metabolism is affected by the loss of TRIP12 (Figure 3 *N*) as well as several features of cell injury responses (Figure 3*O*). Finally, we noticed the increase of expression of several genes coding for nucleic acids binding proteins including chromatin remodeling factors (Figure 3*P*).

Altogether, our results suggest that TRIP12 deficiency during embryogenesis leads to an altered mature acinar cell identity.

### TRIP12 deficiency in adult acinar cells does not alter the acinar cell morphology

Since PDX1-expressing cells give rise to all the pancreatic lineages and adult cell types, we generated a conditional model consisting in an acinar-cell specific ablation of Trip12 gene to study the role of TRIP12 more specifically in these cells and to avoid possible artefacts due to impact on endocrine pancreas. We used Elas-CreER mice in which a conditional disruption of Trip12 gene in adult mice is induced by an administration of tamoxifen. In parallel, we combined an inducible deletion of Trip12 gene to a lineage tracing by using ZsGreen reporter mice (Figure 4*A*). As expected from the Cre-activity pattern in Elas-CreER mice, ZsGreen expression is present in acinar cells but absent in ductal cells and islet cells (Figure 4*B*). The pancreas to body weight ratio of Elas-CreER;Trip12^f/f^ TRIP12-deficient mice (ET) significantly decreases at five months of age, three months after tamoxifen treatment. The same trend, although not statistically different, is observed in older mice (Figure 4*C*). They exhibit no macroscopic modifications of their pancreas and H&E staining shows similar pancreas phenotypes for control and TRIP12-deficient mice (Figure 4*D*). Nevertheless, we confirmed acinar Trip12 deletion by immunofluorescence in ET pancreas using the same previously described anti-TRIP12 antibodies (Figure 4*E*). While TRIP12 is heterogeneously present in the nuclei of murine acinar, ductal and islet cells of control E mice, the loss of TRIP12 is exclusively visualized in acinar cells of ET mice pancreas.

**Figure 4:**
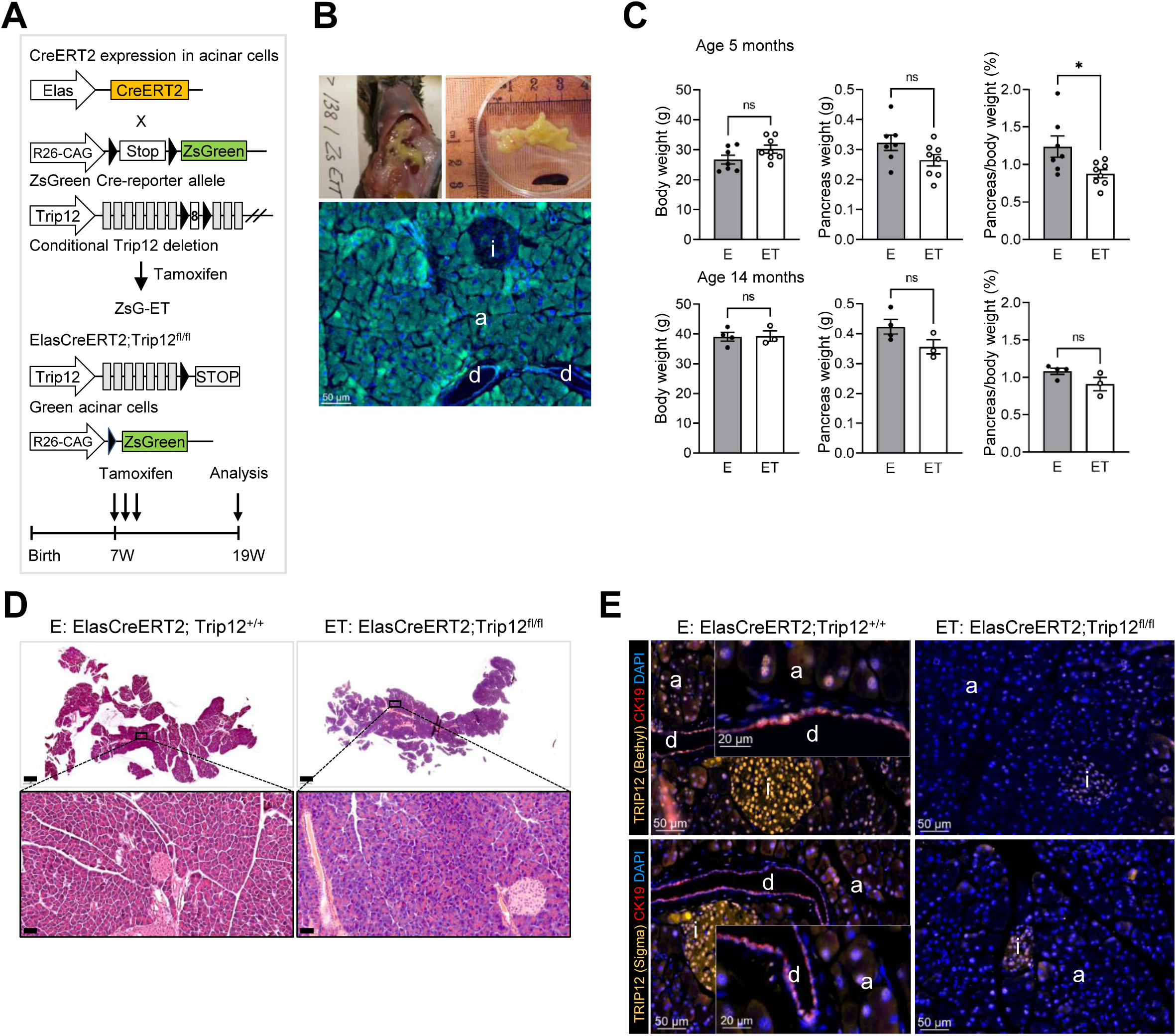
TRIP12 deficiency in adult acinar cells does not the alter acinar cell morphology. ***(A)*** Schematic representation of the ET model and of ZsGreen conditional expression in acinar cells in which the CreER recombinase expression is driven by the Elastase Cre^ERT2^ promoter. Black triangles indicate loxP sequences. Experimental approach is described. ***(B)*** Expression of ZsGreen in the pancreas of ZsG-ET mice showing specific recombination in acinar cells (a). Nuclei are counterstained with DAPI, ductal cells (d), islet cells (i). Scale bar: 50μm. ***(C)*** Body weight ratio, pancreas weight and pancreas/body weight ratio of E control mice (n=7) and TRIP12-depleted ET mice (n=6) at the age of five months and of E (n=4) and ET (n=4) mice at the age of 14 months. Data are expressed as mean ± SEM, *indicates a p value lower than 0.05. ***(D)*** H&E staining of E control and ET pancreas four months after tamoxifen treatment, scale bars 1000 µm. Boxes indicate enlarged regions, scale bars: 50 μm. ***(E)*** Immunofluorescence staining of cytokeratin19 (CK19) and of TRIP12 with Bethyl anti-TRIP12 antibody (upper panels) or with Sigma anti-TRIP12 antibody (lower panels) in E and ET pancreas. Nuclei are counterstained with DAPI. Scale bars: 50 μm. Enlarged boxes, scale bars: 20 µm; a: acini, d: duct, i: islet. Images are representative of three independent experiments.

### TRIP12 is required for ADM formation in vitro

Acinar cells undergo acinar-to-ductal metaplasia (ADM), a reprogramming process that form metaplastic ducts in response to inflammation and/or Kras^G12D^ mutation which precedes the formation of PanIN arising from acinar cells. Given the elevated level of TRIP12 in these early-stage preneoplastic lesions and the repertoire of known TRIP12 functions, we investigated whether it plays a role in the regulation of acinar cell plasticity in response to injury. We used an established 3D acinar cell culture to model the conversion of acinar cells and the lineage tracing with the ZsGreen reporter allele to follow the fate of TRIP12-expressing and TRIP12-deficient cells. Acinar cells were isolated from TRIP12-expressing pancreas of ZsG-E and ZsG-C control mice and from TRIP12-deficient pancreas of ZsG-ET and ZsG-CT mice (Figure 3*A* and 4*A*). ADM was initiated by an addition of epidermal growth factor (EGF). TRIP12-expressing and TRIP12-depleted acinar cells from both mice genotypes remain organized in small acinar cell clusters after one day of culture (Figure 5*A* and *B*).

**Figure 5:**
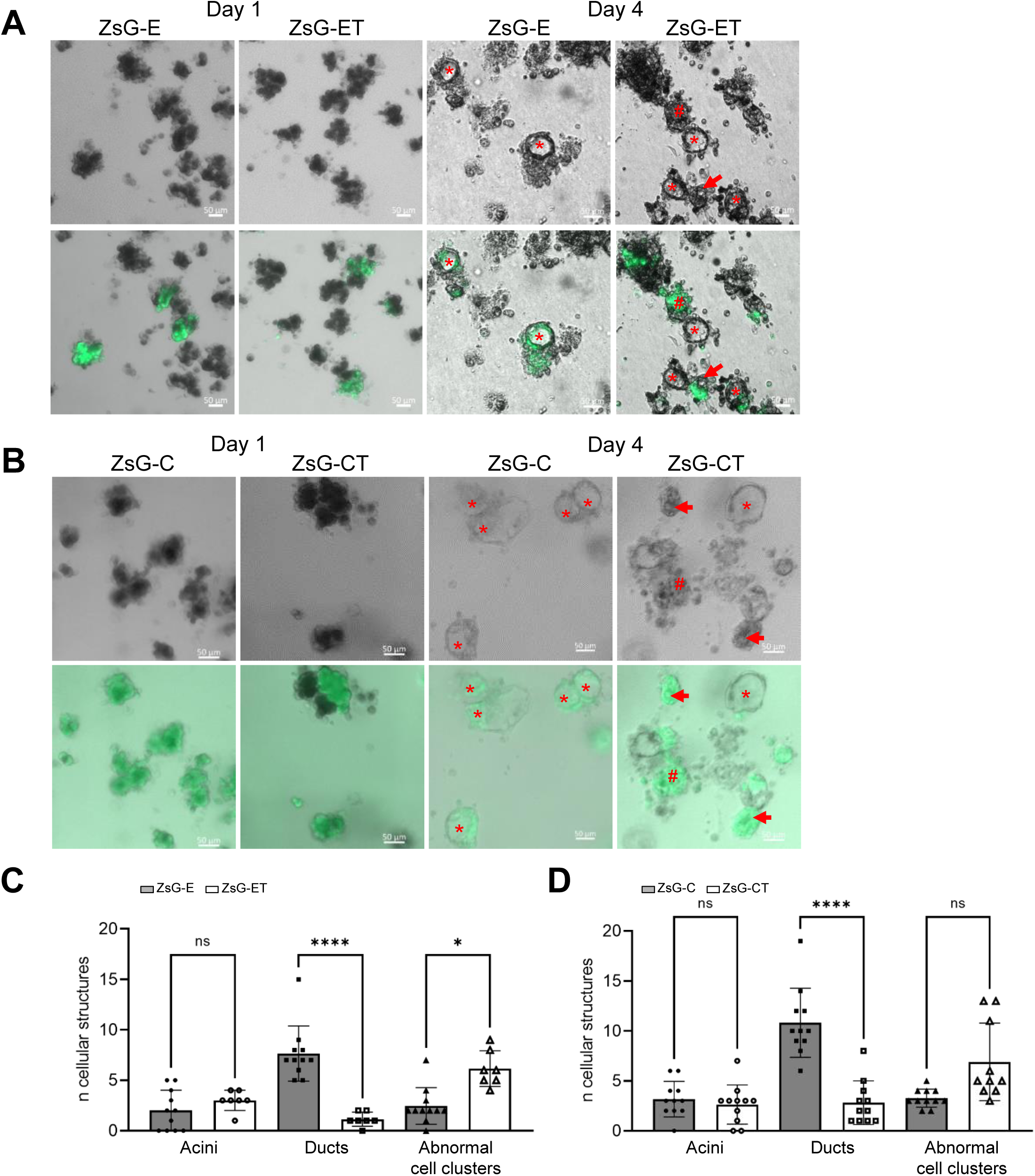
TRIP12 is required for ADM formation in vitro. ***(A-B)*** Bright field and fluorescence images of acinar cell clusters and ductal cyst structures from isolated acinar cells from control ZsG-E and ZsG-C and Trip12-deficient ZsG-ET and ZsG-CT pancreas on day 1 and on day 4 after isolation and treatment with EGF. For these experiments both Trip12^+/+^ and Trip12^fl/fl^ mice were bred with ZsGreen reporter mice prior crossing with ElasCreERT2 mice or Pdx1Cre mice yielding ZsGreen expression in control ZsG-E and ZsG-C and in ZsG-ET and ZsG-CT Trip12-deficient acinar cells. Red asterisks indicate ductal cysts structures forming after ADM. Red hashes indicate fluorescent acinar cell clusters that do not undergo acinar to ductal metaplasia. Red arrows show abnormal ET and CT acinar cell clusters with enlarged lumen or vacuoles but no ADM. Scale bars: 50 μm. Images are representative of three independent experiments. ***(C-D)*** Quantification of fluorescent cellular structures: acinar cell clusters, ductal cyst structures and abnormal acinar cell clusters (characterized by multilobed or enlarged lumen/vacuoles and by an absence of ductal flat epithelium) by day 4 after acinar cell isolation from ZsG-E and ZsG-C control (grey bars) and ZsG-ET and ZsG-CT (white bars) pancreas and treatment with EGF. The bars represent the means ± SD of three independent experiments.

As expected, after four days of culture in the presence of EGF, control acini convert to ductal cyst structures with a single cellular layer surrounding a lumen (Figure 5*A* and *B*). Importantly, ZsGreen expression does not prevent ADM. Unlike control TRIP12-expressing acini, TRIP12-depleted acinar cells do not transdifferentiate into ductal cells. The ductal structures observed in ZsG-ET and ZSG-CT TRIP12-deficient mice never contain ZsGreen-expressing cells. Accordingly, these cells correspond to cells that failed to undergo any Cre-mediated recombination and therefore express TRIP12. The absence of TRIP12 is linked to the presence of fluorescent cell clusters of various sizes where sometimes a tiny lumen or vacuole can be distinguished. In any cases, these abnormal cell clusters never resemble ductal cysts. Quantification of these cell clusters and of metaplastic ducts demonstrates that TRIP12 is required for acinar to ductal metaplasia (Figure 5*C* and *D*) and contributes to acinar cell plasticity.

### TRIP12 is required for ADM formation in vivo

Our 3D-culture model recapitulates acinar cell dedifferentiation similarly as it occurs during pancreatitis. We therefore induced acute pancreatitis in control and TRIP12-depleted mice to investigate the role of TRIP12 in vivo. To avoid any impact on endocrine pancreas and effects on cell injury response linked to the absence of TRIP12 in embryonic pancreas, we used ET mice and analyzed histological changes in response to pancreatic injury at three time points: 2 hours, 1 day and 5 days (Figure 6*A*). Caerulein was injected to mice at the age of 8 weeks. Pancreas of TRIP12-deficient and pancreas of control mice are similar when treated with vehicle (Figure 6*B*). Induction of pancreatitis is confirmed by an increase of plasma amylase level which is similar in both groups (Figure 6*C*). Two hours after the last caerulein injection, pancreas of both genotypes is edematous and infiltrated by immune cells (Figure 6*D*). Vacuolization of acinar cells revealing autophagy and/or stressed acini or necrosis is observed in both control mice and TRIP12-deficient mice pancreas. Distension of acinar lumens related to an initiation of ADM is observed in control mice while the response to injury is different in TRIP12-depleted acinar cells. They exhibit acinar hypertrophy, eosinophilic zymogen granular zones more evenly distributed throughout cells, and decreased basophilic acinar basal regions compared to controls. In control mice, the pancreatic alteration becomes more pronounced as the inflammatory process progresses. An analysis 24 hours after injury displays entire lobules of control mice with extensive ADM where acinar cells have lost polarity and adopt a columnar morphology similar to ductal cells. In contrast, TRIP12-depleted acini do not evolve towards a ductal phenotype. They retain a hypertrophied acinar phenotype and are characterized by the presence of numerous vacuoles of different sizes that contain cellular or amorphous material (Figure 6*D*). While interstitial edema area is similar in pancreas of both genotypes, inflammatory cells are more numerous in control mice pancreas one day after caerulein treatment (Figure 6*E*). Five days after caerulein treatment, most of control mice acinar tissue reverts back to its original morphology whereas ET acinar cells retain several defects. Vacuoles are still present suggesting that longer duration is required for a complete recovery in the absence of TRIP12 (Figure 6*D*).

**Figure 6:**
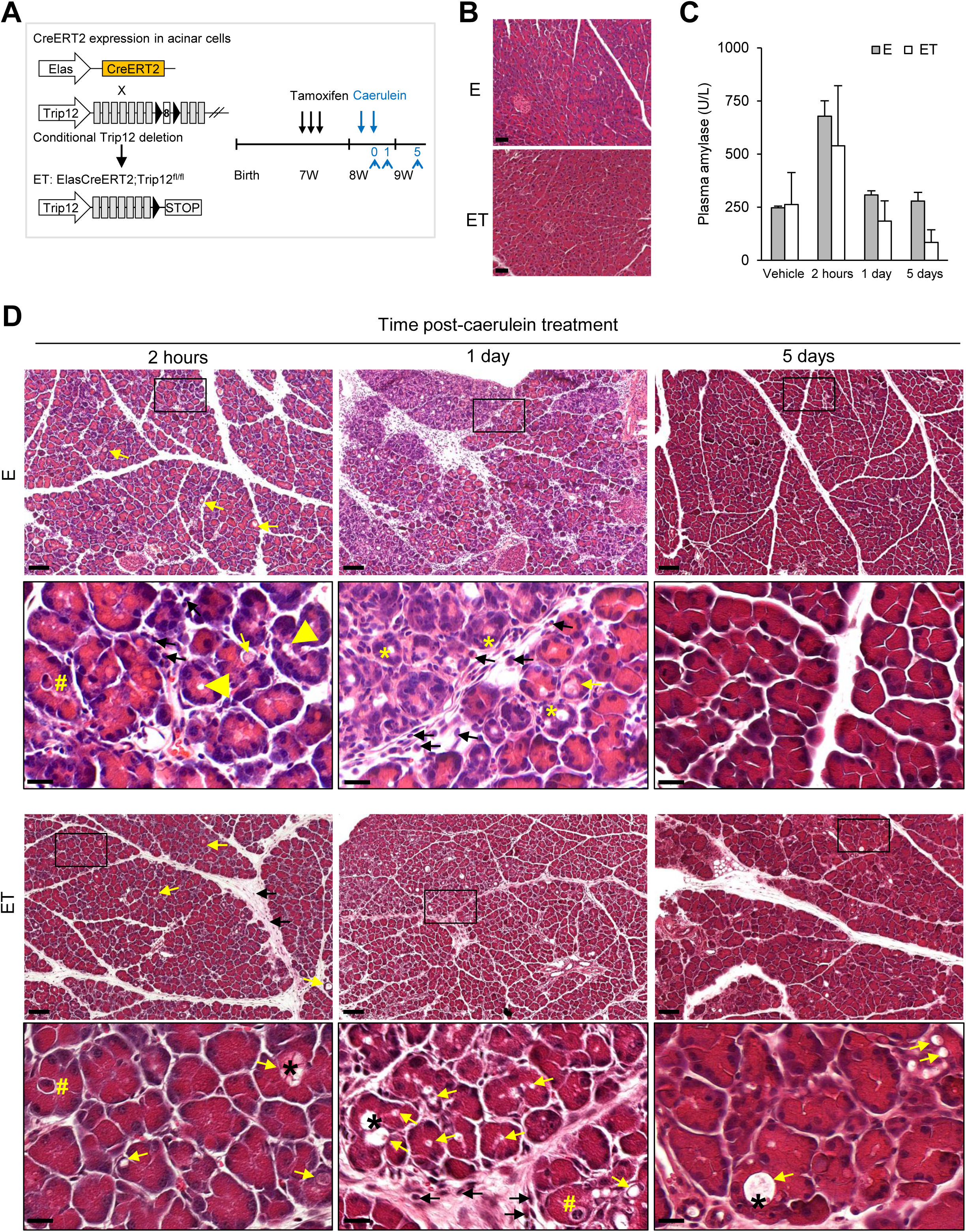
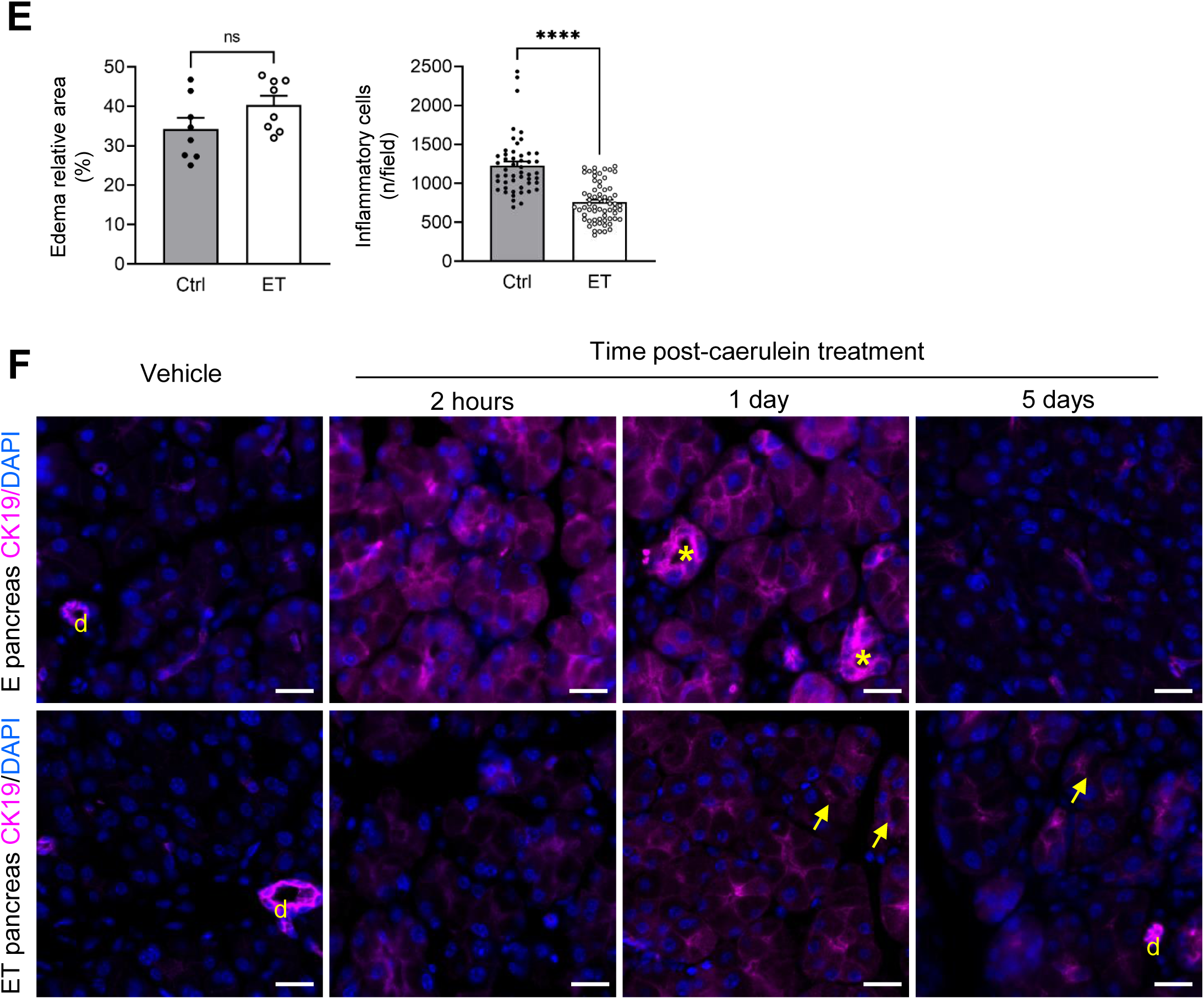
TRIP12 is required for ADM formation in vivo. ***(A)*** Experimental design of caerulein-induced acute pancreatitis in the ET and E control mice. Inactivation of TRIP12 was induced by injections of tamoxifen at the age of seven weeks. Controls were also treated with tamoxifen. Pancreatitis was induced one week later. Mice were sacrificed 2 hours (0), 24 h (1) and 5 days (5) from the last injection. ***(B)*** H&E sections of E and ET mice. The images are representative of 5 ET and 10 E mice treated 24 hours with vehicle, scale bars: 50 µm. ***(C)*** Plasma amylase levels of E (grey bars, n=3) and ET (white bars, n=3) mice were determined at 2h, 1 day or 5 days after the last injection. Results are expressed as a mean ± SEM. ***(D)*** H&E sections of E and ET mice treated with caerulein and analyzed at 2h, 1 day and 5 days after the last injection, scale bars: 100 μm. Enlarged boxes (scale bars: 20 µm) show control acinar cells with central distended lumen at 2 hours (yellow arrowheads) that evolves towards a ductal phenotype by day 1 post-treatment (yellow asterisks). E and ET acini exhibit vacuoles of various sizes (yellow arrows) sometimes containing amorphous material (black asterisks). Yellow hashes indicate apoptotic cells. Inflammatory cells (black arrows) are present in the interstitial edema. Images are representative of three mice at each time. ***(E)*** Quantification of pancreatic injury in E and ET pancreas by day 1 after the last caerulein treatment. Interstitial edema area/pancreas ratio was measured in two sections and inflammatory cells were counted in 10 to 25 20x-fields of three E and three ET mice pancreas using Fiji software. Data are expressed as means ± SEM; ns, non significant, ****p<0.0001. ***(F)*** CK19 expression at the indicated times after caerulein treatment. Nuclei are counterstained with DAPI. Scale bars: 20μm, d: duct. Yellow asterisks indicate transdifferentiating acini, yellow arrows show vacuoles. Images are representative of three experiments with three different mice.

In accordance with ADM, the pancreas of control mice displays an increased CK19 staining in acinar cells compared to an absence of expression in acinar cells of vehicle-treated mice. CK19 expression is observed from two hours, peaks by day 1 and returns to basal levels at day 5 (Figure 6*F*). CK19 expression is also detected two hours after caerulein treatment in TRIP12-depleted acini but at lower levels than in control mice at the same time point. CK19 expression slightly increases by day 1 in the absence of TRIP12 confirming the altered differentiation response to caerulein. Unlike control pancreas, CK19 immunostaining is restricted to centroacinar lumens and does not extend in basolateral location as observed in ADM (Figure 6*F*).

Our experiments demonstrate that TRIP12 is necessary for ADM formation in vivo and its depletion alters differentiation response to caerulein in mice pancreas.

### TRIP12 cooperates with Kras^G12D^ to initiate pancreatic carcinogenesis

Several studies showed that Kras^G12D^ oncogene expression in adult acinar cells initiates PDAC.^38^ We hypothesized that altered ADM that is observed in absence of TRIP12 has consequences on PDAC initiation. To investigate the role of TRIP12 in pancreatic carcinogenesis, we crossed Elas-CreER;Trip12^f/f^ mice with LSL-Kras^G12D^ mice to generate TRIP12-deficient KET and TRIP12-expressing KE mice (Figure 7*A*). The activation of Kras^G12D^ and the loss of TRIP12 were achieved by an administration of tamoxifen. Mice were sacrificed three months after tamoxifen treatment. We observed no macroscopic changes between KE and KET mice and their pancreas in both conditions (Figure 7*B*). An histological analysis reveals the presence of low grade PanIN lesions in the pancreas of KE mice but no evidence of preneoplastic lesions in any of the KET mice pancreas that exhibit normal acinar tissue (Figure 7*D*).

**Figure 7:**
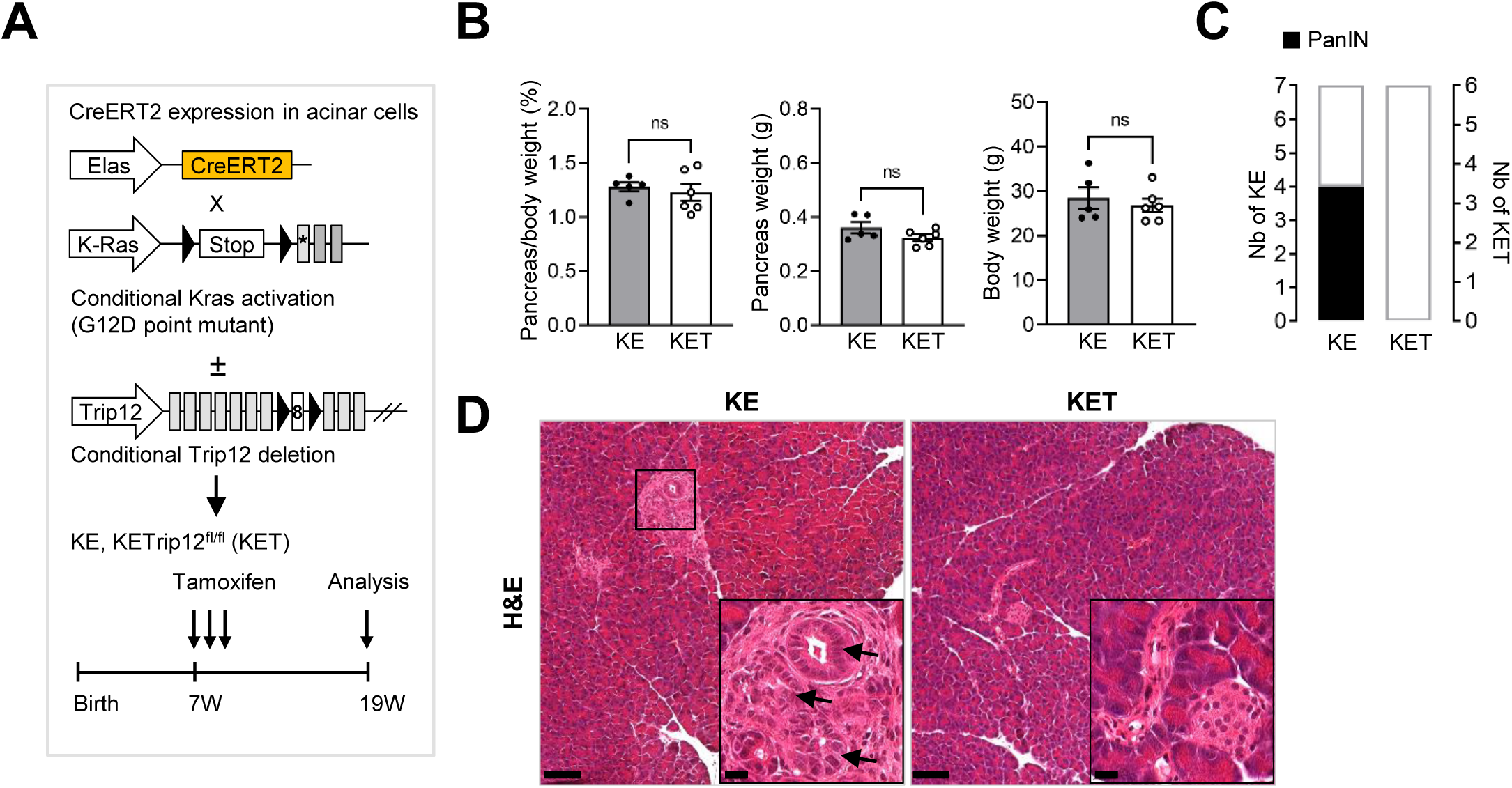
TRIP12 cooperates with Kras^G12D^ to initiate pancreatic carcinogenesis. ***(A)*** Schematic representation of the KET murine model. Black triangles indicate loxP sequences. TRIP12 deletion and KrasG12D activation are induced by tamoxifen at the age of seven weeks, mice are sacrificed at the age of 19 weeks. **(*B*)** Pancreas/body weight ratio, pancreas and mouse weight of KE (n=5) and KET mice (n=6) three months after tamoxifen treatment. Data are presented as means ± SEM, ns, non significant. ***(C)*** Incidence of precancerous PanIN lesions in KE and KET mice. ***(D)*** Histological analysis of KE and KET mice, scale bars 100 µm, boxes indicate enlarged regions in insets, scale bars: 20 μm. Black arrows indicate PanIN lesions. Images are representative of seven KE and six KET mice.

These results demonstrate that an inactivation of TRIP12 prevents the oncogenic Kras-induced PanIN formation. They indicate that TRIP12 is a factor necessary for the initiation steps of pancreatic carcinogenesis. They also suggest that TRIP12 regulation of acinar cell plasticity is required for transformation by Kras^G12D^ oncogene.

### TRIP12 depletion limits tumor cell metastatic capacity

TRIP12 is overexpressed in pancreatic cancers (Figure 1*F* and *K*). Kaplan-Meier analysis of the patient survival stratified by expression level of TRIP12 shows that patients with a lower TRIP12 expression exhibit a better survival than patients with a higher TRIP12 expression (Figure 1*D*). We previously demonstrated that TRIP12 exerts important functions during cell cycle and that a depletion of TRIP12 significantly inhibits cancer cell growth.^35^ We hypothesized that TRIP12 influences PDAC growth and progression. We generated mice cohorts of the consensus KPC mouse PDAC model combining concomitant endogenous expression of Kras^G12D^ and Trp53^R172H^ with (KPCT mice) or without (KPC mice) TRIP12 ablation (Figure 8*A*). Autopsy and histological analysis reveal that both KPC and KPCT mice develop pancreatic ductal adenocarcinoma (Figure 8*B*). Clinical spectrum of the disease shows similar body and tumor weights of KPC and KPCT (Figure 8*C*) but a higher tumor incidence and more frequent metastasis in KPC than in KPCT mice (Figure 8*D*). Median survival of KPCT mice is not significantly increased compared to that of KPC but a delay in the death of mice with the less aggressive KPCT tumors is observed (Figure 8*E*).

**Figure 8:**
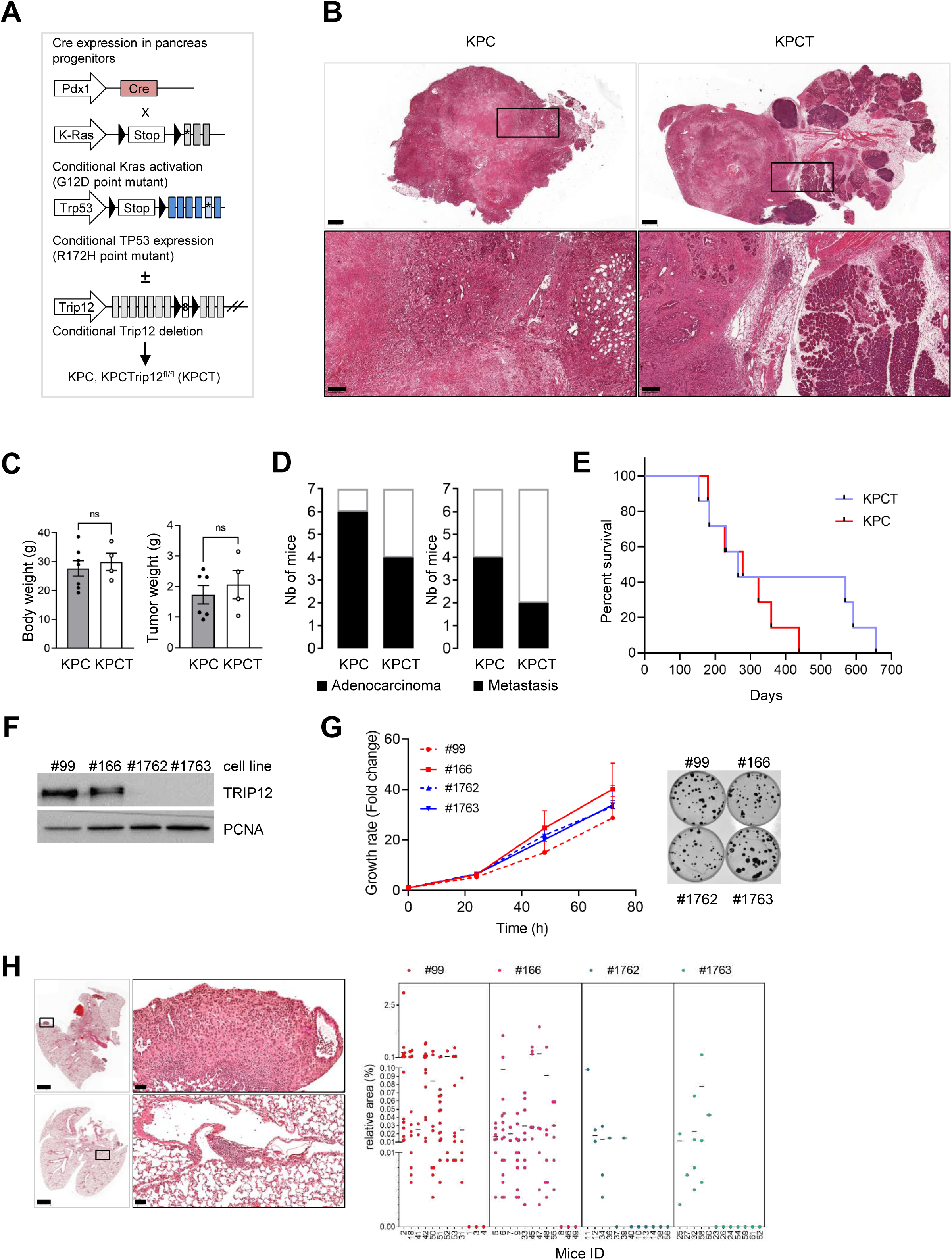
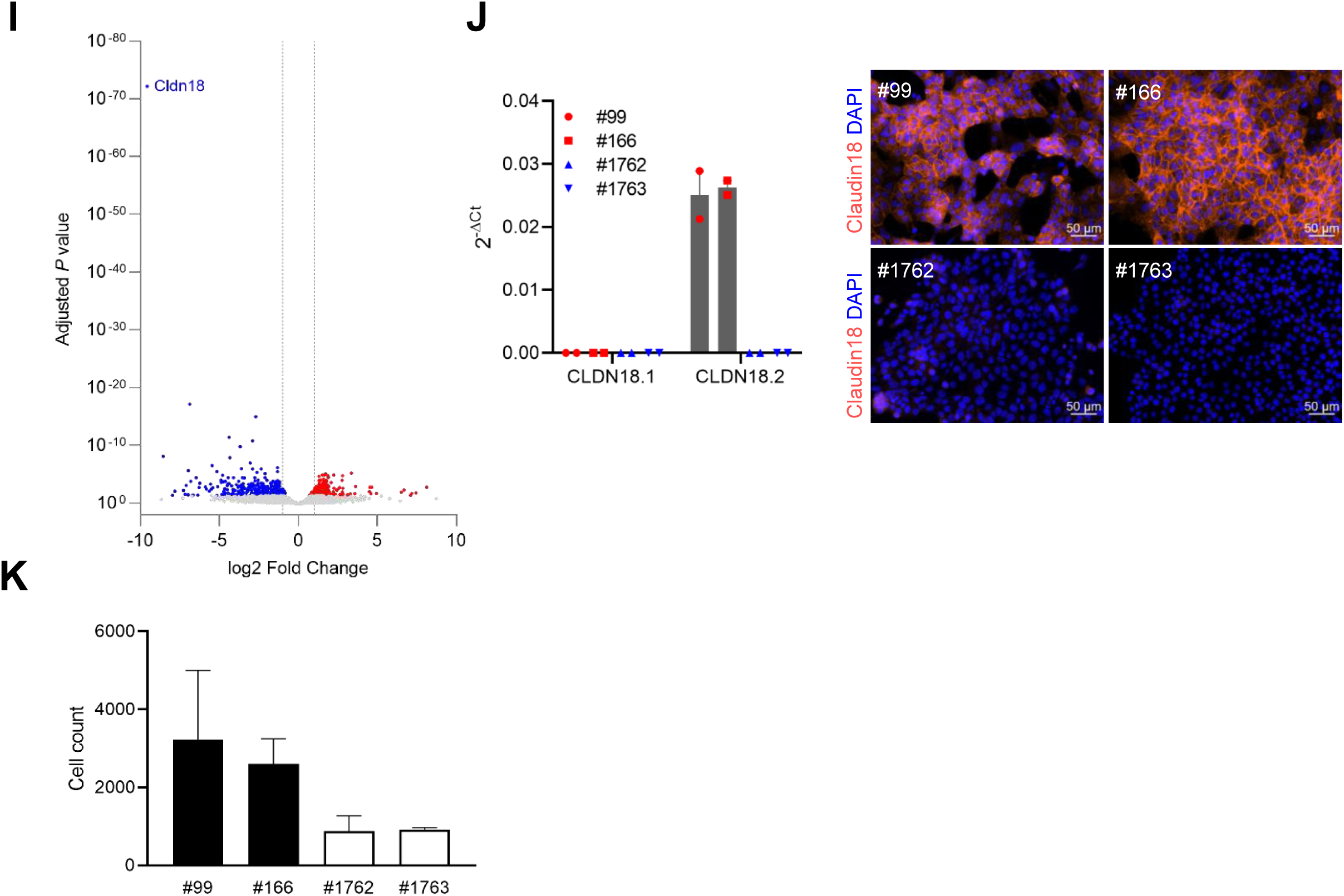
TRIP12 depletion limits tumor cells metastatic capacity. ***(A)*** Schematic representation of the KPC and KPCT models. Black triangles indicate loxP sequences. ***(B)*** Representative H&E staining of KPC and KPCT pancreas, scale bars: 1000µm. Black boxes indicate enlarged regions (scale bars: 200 µm). ***(C)*** Body weight of KPC (n=7) and KPCT (n=4) mice, tumor weight in KPC (n=6) and KPCT (n=4) pancreas. ***(D)*** Incidence of adenocarcinoma and metastasis in KPC (n=7) and KPCT (n=7) mice. The number of mice with adenocarcinoma and metastasis is represented with black bars. ***(E)*** Survival of KPC (n=7, red curve) and KPCT (n=7, blue curve) mice. ***(F)*** TRIP12 protein expression in KPC and homozygous KPCT cell lines derived from pancreatic tumors of KPC and KPCT mice was determined by a Western-blot analysis. The image is representative of at least three separate experiments. PCNA is used as a loading control ***(G)*** Left panel: cell growth rate of KPC and KPCT derived murine cell lines determined by cell counting, results are expressed as mean ±SEM compared to t=0h (set as 1) of at least three independent experiments. Right panel: colony formation assay of KPC and KPCT cells, colonies are stained using Crystal violet, n=3. ***(H)*** Left panel: representative H&E staining of lung metastasis from C57Bl6 mice that were injected in the tail vein with cells derived from pancreatic tumors of KPC and KPCT mice (scale bars: 2000 µm). Black boxes indicate enlarged regions (scale bars: 50 µm). Right panel: Relative area of metastasis area in lungs of C57Bl6 mice that were injected in the tail vein with cells derived from pancreatic tumors of KPC mice. Twelve mice were injected with each of the cell lines (KPC #99 and #166, KPCT #1762 and #1763). The surface of the metastasis was quantified on two whole lung sections for each mouse and normalized with the total area of the lungs. Results are expressed as the relative area of all metastasis for each mouse. Mouse ID are indicated on the x axis. ***(I)*** Volcano plot of genes with differential mRNA expression in KPCT cells from RNA-Seq of three biological replicates for KPC and KPCT cell lines. Blue dots represent genes with a reduced expression, red dots represent genes with an increased expression, grey dots represent genes with an adjusted P value lower than 0.05. **(*J)*** Left panel: qRT-PCR analysis of Claudin 18 isoforms (CLDN18.1 and CLDN18.2) in TRIP12-expressing cells #99 and #166 (red symbols) and in TRIP12-deficient cells #1762 and #1763 (blue symbols). Data are representative of three independent experiments. Right panel: Claudin 18 expression (orange) in the indicated tumor cell lines, nuclei are counterstained with DAPI (blue). Scale bars: 50μm ***(K)*** Invasion assay of #99 and #166 cells (black bars) and of #1762 and #1763 cells (white bars). Results are expressed as the mean ±SEM (n=3).

We generated pancreatic cancer cell lines from tumors of KPC (#99 and #166) and from tumors of KPCT (#1762 and #1763) mice (Figure 8*F*). Proliferation and colony formation assays show no statistical differences between KPC and KPCT cell lines (Figure 8*G*). To model metastasis formation, we carried out tail vein injections of these KPC cancer cells lines in C57/B6 mice. The results of these experiments demonstrate that the depletion of TRIP12 inhibits in vivo metastatic dissemination. Metastatic foci are more numerous and their area larger after tail vein injection of syngeneic KPC pancreatic cells compared to syngeneic KPCT pancreatic cells (Figure 8*H*). We subjected total RNA extracted from KPC and KPCT cell lines to a RNA-Seq analysis. A total of 609 genes are significantly altered (adjusted P value ≤ 0.05), (Figure 8*I*). We noticed that the most downregulated gene in KPC cells in the absence of TRIP12 encodes for the tight junction protein Claudin-18 (adjusted P value = 6.57.10^-73^) (Figure 8*I*). We assessed the expression of the two CLDN18 transcripts (CLDN18.1 and CLDN18.2) in KPC and KPCT cells by RT-qPCR. As expected, no transcript of the lung specific isoform CLDN18.1 is detected in both KPC and KPCT murine pancreatic tumor cell lines (Figure 8*J*). KPC cells express CLDN18.2 transcripts but they are absent in KPCT cells (Figure 8*J*). We also confirm the RNA data at the protein level by immunofluorescence (Figure 8*J*). We then used Boyden chambers to determine if the loss of TRIP12 affects cell invasion in KPCT cells and found that cell invasion decreases in KPCT cells (Figure 8*K*).

These findings indicate that TRIP12 facilitates pancreatic tumor formation, tumor cells invasion and likely regulates Claudin-18 expression.

## Discussion

In the present study, we provide convergent lines of evidence supporting a new key role of TRIP12 both in the induction of ADM and the formation of PanIN and in early metastatic dissemination. We demonstrate that TRIP12, which is expressed at a low level in the healthy pancreas, is markedly overexpressed in pancreatic preneoplastic lesions and pancreatic cancers. Using two conditional Trip12 knockout mice models, we reveal that TRIP12 expression is required for ADM formation. We also show that TRIP12 deficiency impedes the formation of PanIN in the presence of mutant Kras^G12D^ and reduces the metastatic capacity of tumors cells when mutated Trp53^R172H^ is concomitantly present.

TRIP12 deletion during embryogenesis has a noticeable impact in adults causing pancreatic steatosis associated with reduced pancreas weight that may reflect impaired pancreatic exocrine function. Deficiency of UBR1 (Ubiquitin Protein Ligase E3 Component N-Recognin 1), an E3 ubiquitin ligase of the N-end rule pathway^39^, induces similar defects in the exocrine pancreas of mice and in Johansson-Blizzard Syndrome patients.^40, 41^ Although TRIP12 deficiency in pancreatic progenitors does not reveal major pancreatic abnormalities in embryos, an analysis of acinar cell RNA population by RNA-Seq shows that their acinar cell identity is altered in adult pancreas without histological changes. The trimeric transcription factor Pancreas Transcription Factor 1 complex (PTF1), formed by the bHLH (beta helix loop helix) transcription factor PTF1a with recombining binding protein suppressor of hairless-like (RBPJ-L) and another common bHLH E-protein, is central in maintaining the differentiation and function of acinar cells.^42^ We find that PTF1a mRNA expression remains unchanged following TRIP12 invalidation. This supports our previous demonstration of a post-translational regulation of PTF1a via TRIP12-mediated ubiquitination^34^ that does not operate in TRIP12-depleted acinar cells and leads to increased protein levels. This, with increased RNA levels of RBPJ-L in TRIP12 knockout acinar cells, likely maintains the active form of PTF1 complex. The RNA level of other key transcription factors genes regulating acinar cell differentiation and organization (i.e., NR5A2, GATA6, and MIST1) and of classical ductal markers (i.e., SOX9 and cytokeratin 19) are unaffected by a loss of TRIP12. Unexpectedly, progenitor genes such as nestin and several Notch pathway genes are induced in the absence of TRIP12 further supporting a change of acinar differentiation or a blockage in a selective acinar cell stage intermediate to ductal differentiation. Both nestin and Notch pathway genes are present during pancreas development and expressed during transdifferentiation of acinar cells into ductal cells^43^. In adult acinar cells, Notch signaling contributes to a recovery from pancreatic injury^44^ and an expansion of an undifferentiated precursor population^45–48^. This indicates that the expression of transcriptional networks main actors operating in differentiated acinar cells is preserved in TRIP12-deleted acinar cells. This also suggests that other mechanisms leading to a downregulation of specific acinar genes and reexpression of developmental genes are involved to modify acinar cell identity.

Our findings highlight the key role of TRIP12 in acinar cell identity and plasticity. They show that a loss of a single protein blocks the reprogramming process in response to pancreatic injury without the influence of other cell types. Of importance, TRIP12 deficiency also impedes the formation of ADM in presence of mutant Kras^G12D^ indicating that TRIP12 is required for the transformation by the Kras oncogene. Increased expression of PTF1a and RBPJ-L in Trip12 knockout acinar cells in physiological conditions could explain the resistance of acinar cells to ADM and PDAC initiation. Loss of PTF1a is already well demonstrated to sensitize acinar cells to ADM, to increase their susceptibility to PDAC initiation and to activate the endoplasmic reticulum stress pathway.^28, 49^ The downregulation of PTF1a is a cell-autonomous event necessary to initiate PanIN formation and a sustained expression of PTF1a prevents Kras-mediated oncogenesis.^31, 32^ This strongly supports our findings that identify TRIP12 as an essential actor of acinar cell differentiation.

The role of epigenetic regulation in the initiation steps of PDAC is now demonstrated.^50^ Transient expression of reprogramming factors (i.e., OCT3/4, SOX2, KLF4, c-MYC)^51^ has been demonstrated to induce ADM.^17^ The repression of acinar cell enhancers that reduces the expression of acinar cell transcription factors and increases expression of ductal genes is now recognized as an ADM inducer. Coherently, deletion of acinar-cell transcription factors leads to ADM.^27, 28, 52–54^ Inversely, a forced expression of these factors prevents the ADM formation.^31, 32^ During ADM, the epigenetic repression of acinar enzymes and transcription factors is initiated and maintained by the repressive histone marks H2AK119Ub (monoubiquitination of histone H2A on lysine119) and H3K27me3 (trimethylation of histone H3 on lysine27) catalyzed by histone modifying enzymes belonging to the polycomb repressive complexes (PRC) 1 and 2, respectively.^19, 20^ Several evidence from the literature support the fact that TRIP12 ubiquitin ligase activity could contribute to repress acinar cell enhancers and that a TRIP12 deletion could inhibit a repressive pathways in acinar cells during ADM. Indeed, we showed the tight interaction of TRIP12 with the chromatin.^35^ Moreover, several TRIP12 protein partners regulate chromatin remodeling complexes^33^ and TRIP12 was demonstrated to control the integrity of the SWI/SNF chromatin remodeling complex.^55^ Furthermore, TRIP12 also mediates the proteolysis of ASXL1, a component of the Polycomb Repressive Deubiquitinase (PR-DUB) that deubiquitinates histone H2AK119.^56^ Therefore, by preserving polycomb gene silencing, TRIP12 might contribute to the repression of acinar genes. An ablation of TRIP12 would then prevent the epigenetic reprogramming of acinar cells and maintain a differentiated acinar state that is sufficient to counteract local inflammation and cellular injury response and to resist to oncogenic Kras effects. Strikingly, a loss of RING1b, the PRC1 E3 ubiquitin ligase catalyzing the ubiquitination of H2AK119 impairs caerulein-induced ADM in vivo and in vitro and decreases the vulnerability to Kras transformation by maintaining acinar cells differentiated state.^20^ A similar phenotype is also observed with a conditional loss of BMI1, another PRC1 component.^57^ Our study revealing a preservation of PTF1a and an increase of RBPJ-L in TRIP12 knockout acinar cells in physiological conditions might corroborate the proposed repressive role of TRIP12 during ADM acting by ubiquitinating PTF1a for proteasomal degradation and limiting chromatin accessibility for RBPJ-L transcription. Given the effects of TRIP12 deletion on inflammation, ADM and transformation (this study), it will be important to determine which and how genes are directly suppressed or activated by TRIP12.

Our results show that whereas it is required for cancer initiation, TRIP12 is dispensable for pancreatic cancer when the oncogenic Kras^G12D^ is associated with the loss of a tumor suppressor. Interestingly, mice survival curves have a pattern similar to that of human patients. Although the median survival is unaffected by TRIP12 expression, patients with low TRIP12 expressing-tumours and TRIP12-knockout KPC mice live longer. We also show that whereas the absence of TRIP12 does not suppress tumorigenic potential in the presence of mutated Kras^G12D^ and Trp53^R172H^, TRIP12 is needed for tumor cells invasive capacity. Previous data from the literature revealed an EMT inhibitory role for TRIP12.^58^ We found no impact of TRIP12 on the epithelial most common character of KPC tumor cell lines, but our data demonstrate that TRIP12 is necessary to switch on the expression of the tight junction protein CLDN18.2 in murine pancreatic KPC tumor-derived cell lines. Two CLDN18 isoform transcripts are produced by an alternative splicing.^59^ In normal tissues the expression of CLDN18.1 is lung specific whereas CLDN18.2 is restricted to the stomach. The CLDN gene expression is frequently altered in various cancers^60, 61^ and CLDN18.2 is aberrantly expressed in several cancers. In PDAC, CLDN18.2 appears in precancerous lesions^62–65^ and an increased expression correlates with the presence of distant metastasis.^66^ To date, mechanisms governing CLDN18 expression are partially known. The silencing of CLDN18.2 in the vast majority of normal tissues has been related to the methylation of its gene promoter^67, 68^ and transcriptional activation linked to the transcription factor CREB in HEK293 cells^67^, to specific protein kinase C signaling pathways in human pancreatic cells^68^ and to the PKC/MAPK/AP-1 signaling pathway^69^ in gastric cancer cells. TRIP12 probably contributes to the regulation of CLDN18.2 expression by a mechanism that remains to be studied.

Finally, the proteasome activity is required for the formation of PanIN.^70^ To our knowledge, the role of HECT E3 ubiquitin ligases has been poorly investigated in pancreatic cancer initiation. Our findings identify TRIP12 as an essential player in acinar-to-ductal reprogramming, in Kras-driven PanIN formation and in tumoral invasion. Otherwise, our data may support strategies aimed at preserving or reexpressing the acinar differentiation. The reintroduction of PTF1a in the context of Kras signaling in pancreatic premalignant cells or cancer cells rescues the acinar gene program and decreases cancer-associated properties. ^31, 32^ Therefore our data suggest that the targeting of overexpressed TRIP12 would contribute to inhibit Kras-driven tumorigenesis. Our data also support the targeting of CLDN18.2 with the tumor cell-selective therapeutic antibody zolbetuximab currently in clinical testing.^66, 71^ Evaluation of TRIP12 status could provide a complementary stratification marker for this therapy.

## Acknowledgements

We thank F. Fiore for CIPHE service monitoring, all members of animal facility of CREFRE, Imag’IN platform and pathology department of Institut Universitaire du Cancer de Toulouse, C. Segura from the histology platform of Institut des Maladies Métaboliques et Cardiovasculaires de Toulouse. We thank Drs J. Rouquette (RESTORE Research Center), F. Larminat (Institut de Pharmacologie et de Biologie Structurale of Toulouse) and L. Bartholin (Centre de Recherche en Cancérologie de Lyon) for helpfull discussions.

## Methods

### Mice – study approval – genotyping - treatments

To generate conditional Trip12 knockout mice, embryonic stem (ES) cell clones carrying the targeted allele (Trip12^tm1a(KOMP)Wtsi^) were obtained from the KnockOut Mouse Project (KOMP) repository. Targeted ES cell clones (agouti C57BL/6 parental cell line JM8A3.N1) were microinjected into blastocysts of C57BL/6 (CIPHE, Centre d’Immunophénomique, Marseille, France) to generate mouse line Tm1a which was crossed with CIPHE Flippase mouse line to generate Trip12 ^tm1c Ciphe^ mice carrying the conditional floxed allele.

Mice were housed under specific pathogen-free conditions at the CREFRE (Centre Régional d’Exploration Fonctionnelle et de Ressources Expérimentales) animal facility (Toulouse, France). All animal studies were approved by the Ministère de l’Education Nationale, de l’Enseignement Supérieur et de la Recherche (APAFIS#3600-201512160838611v3) and performed in accordance with institutional guidelines. The following mouse strains were previously described: the LSL-Kras^G12D^ and LSL-Trp53^R172H^ knock-in mice were obtained from the Mouse Models of Human Cancers Consortium repository (NCI-Frederick, USA), Pdx1-Cre mice were obtained from DA Melton (Harvard University, Cambridge, MA, USA), Elastase Cre^ERT2^ mice (Elas-CreERT2) were generated by DA Stoffers (Philadelphia, PA, USA) and were kindly provided by Pr P Jacquemin (UCL, Belgium). ZSGreen mice (B6.Cg-Gt(ROSA)26Sor^tm6(CAG–ZsGreen1)Hze^/J) were obtained from the Jackson Laboratory. Mice were interbred on mixed background (CD1/SV129/C57Bl6) to obtain Pdx1Cre;Trip12^f/f^ (CT mice), Elas-CreERT2;Trip12^f/f^ (ET mice), LSL-Kras^G12D^;Trip12^f/f^;Elas-CreERT2 (KET mice) and LSL-Kras^G12D^;LSL-Trp53^R172H^;Trip12^f/f^;Pdx1-Cre; (KPCT mice). To perform lineage tracing experiments, we generated ZsGreenCT (ZsG-CT) and ZsGreenET (ZsG-ET) mice. Control mice of CT, ET, KET, KPCT, ZsG-CT and ZsG-ET lack the Trip12 floxed allele and are hereafter referred as C, E, KE, KPC, ZsG-C and ZSG-E respectively.

For genotyping, we used PCR protocols recommended by the NCI for mutant Kras^G12D^ and Trp53^R172H^, by the Jackson Laboratory for ZsGreen detection and by the CIPHE for the detection of Trip12 tm1c allele. Primers sequences used for genotyping are listed in Table 1.

**Table 1.**
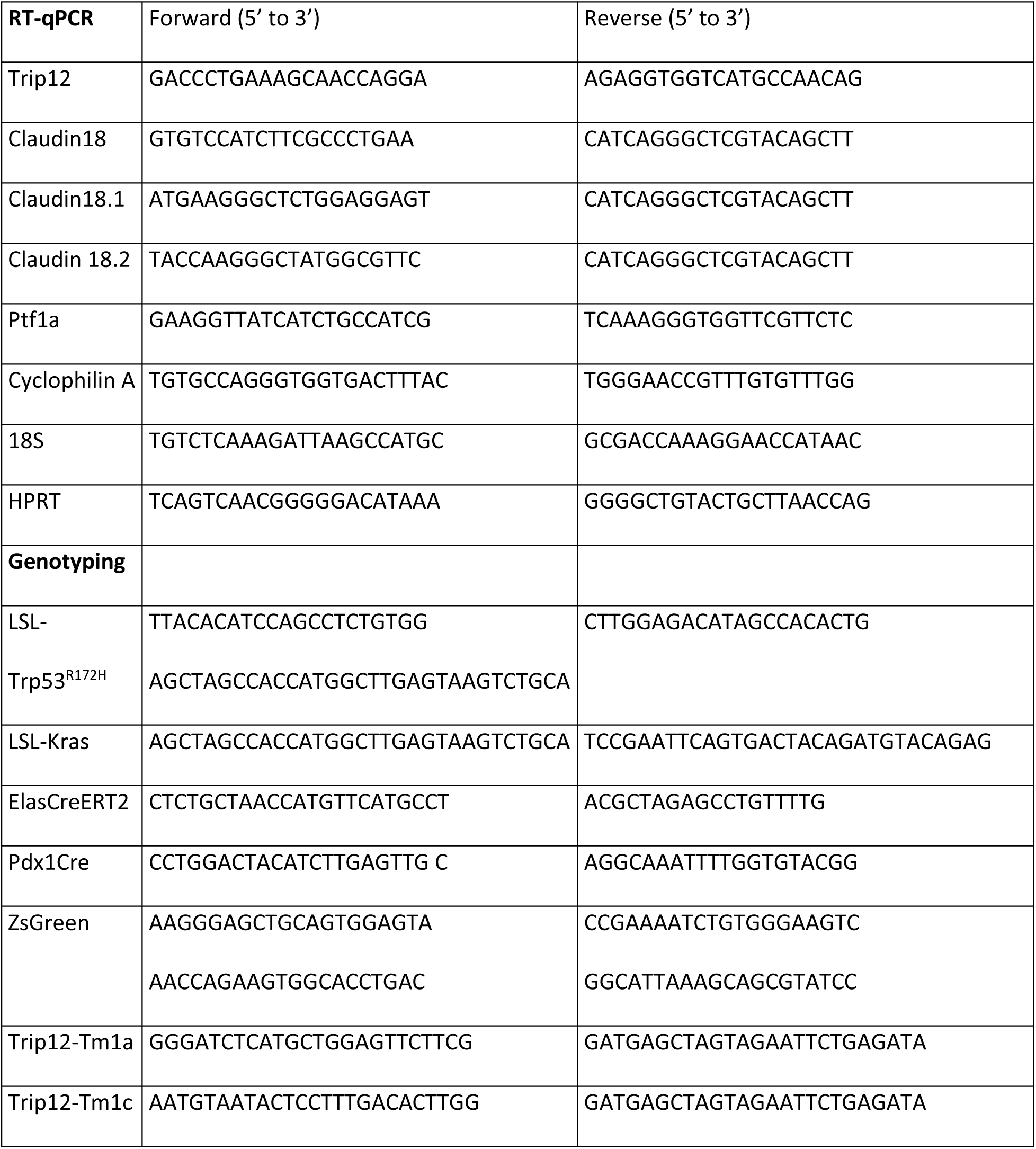
Primers sequences

Seven week-old ET and control mice were treated with tamoxifen (Sigma-Aldrich) dissolved in corn oil containing 10% ethanol at a concentration of 20 mg/ml. Mice received three intra-peritoneal (i.p.) injections (1 mg/15 g of body weight) on three consecutive days.^72^

Acute pancreatitis was induced at 8 weeks of age by two sets of 6 hourly i.p. caerulein injections (50 μg/kg; Sigma-Aldrich) on alternating days separated by 24 hours after a fasting period of 12 hours.^73^ Mice were sacrificed 2, 24 and 120 hours from the end of the experiment. Blood and tissue samples were collected for further analyses. Amylase concentration was measured in plasma using the quantitative assay of alpha-amylase (Phadebas amylase test). Glycemia was determined from tail blood using a glucose monitor (Accu-Chek Performa). Glucose (2 g/kg of body weight) was injected i.p. and glycemia measured at the indicated times.

### Clinical samples

Surgically resected specimens from 27 patients were obtained from the department of pathology (Pr. J Selves, Cancer University Institute of Toulouse) following ethical procedures. They were immunostained with anti-TRIP12 (1:1000, Bethyl Laboratories A301) using standard procedures.

Human tissue samples containing intraductal papillary mucinous neoplastic (IPMN) lesions were provided by the Pathology Department of Beaujon Hospital (Clichy, France). A total of 77 paraffin-embedded IPMN were included in Tissue Micro Array blocks. They were immunostained with an anti-TRIP12 (1:1000, Bethyl Laboratories #A301) using standard procedures.

### Embryos – optical clearing-segmentation

Mice were bred to obtain Pdx1-Cre;Trip12^f/f^ (CT) and control embryos. The day of vaginal plug is defined as E0.5. Adult mice were heavily anesthetized with an overdose of isofluorane. E13.5 and E15.5 embryos were collected in ice-cold phosphate buffer saline, one piece of tail was collected for further DNA extraction and genotyping. Embryos were fixed overnight at 4°C in 10% neutral buffered formalin and again one hour at room temperature the next day. Fixed samples were washed in PBS for one hour and stored at 4°C. For optical clearing, fixed embryos were rinsed with PBS, included in 1% low-melting agarose (Life Technologies) and dehydrated with 100% methanol. They were cleared with the benzyl alcohol/benzyl benzoate (BABB) technique as previously described.^74^ Briefly, embryos were first immersed in a 3/1/2 (v/v) methanol/benzyl alcohol/benzyl benzoate (BABB) solution during 30 min then cleared in 1/2 (v/v) benzyl alcohol/benzyl benzoate. Acquisitions were performed with selective plane illumination microscope (SPIM) technology. MacroSPIM microscope was used for acquisition at 561 nm. Autofluorescence was detected with a 609/57 nm filter and the exposure time was 400 ms. Segmentation of embryos was carried out using Amira software with a dual thresholding method to obtain the volume of the samples. Segmentation of pancreas and liver were done after cropping with Fiji software and using either AMIRA software or manual selection with appropriate thresholding.

### Acinar cell isolation - in vitro acinar to ductal metaplasia formation

Mice were anesthetized with isofluorane and sacrificed by cervical dislocation. Pancreas were quickly resected, rinsed and 100 mg of the organ were placed in 5 ml of cold HBSS (Hank’s balanced salt solution) containing 0.01% (w/v) STI (Soybean Trypsin Inhibitor, Sigma) and 1000 U of collagenase II (Thermofisher). Digestion was performed at 37°C during 20 minutes and mechanical dissociation every 5 minutes through 10 back-and-forth passages of the pancreatic tissue into sterile serological pipettes of decreasing size (25, 10 and 5 ml). Following multiple washes with HBSS supplemented with 5% FCS, 0.01% STI and 10 mM Hepes (pH 7.5), digested pancreatic tissue was pelleted by 400 rpm centrifugation, resuspended in DMEM culture medium containing 4.5 g/l glucose, 10% fetal calf serum (FCS) (Eurobio), 10% penicillin-streptomycin mixture, 0.01% STI and filtered through 100 µm-Nylon mesh. EGF (final concentration: 20 ng/ml) was added to the acinar suspension. An equal volume of neutralized rat tail collagen (RTC) type I (Corning) was added to the acinar cell suspension and 500µl pipetted into wells of 24-well plate precoated with 100 µl of neutralized RTC. Plates were placed at 37% and 5% CO2. After solidification of the RTC, 1 ml of DMEM culture medium (with above mentioned constituents) was added in each well. Medium was changed at day 1 and day 3. Acinar to ductal metaplasia was quantified using Fiji software.

### Generation of mouse cancer cell lines

Tumor cells were isolated using the explant culture method. Briefly, the resected pancreatic tumors from KPC and KPCT mice were cut into small pieces (0.5-1 mm^3^). The tissue pieces were rinsed in DMEM-F12 medium (Sigma) supplemented with penicillin (100 U/ml)/streptomycin (100 µg/ml) and transferred into wells of 6-well plates (4-6 pieces per well) containing 800 µl of culture medium (DMEM-F12 supplemented with 10% FCS and penicillin/streptomycin). The media were renewed every three days and the primary cultures were monitored for 15-35 days for epithelial cell outgrowth. The primary outgrowing epithelial cells were subcultured by trypsinization using a 0.05% trypsin - 0.02% EDTA solution (Thermofischer) and plated into 25-cm^2^ flasks in DMEM-F12 supplemented with 10% fetal calf serum and penicillin/streptomycin. From the second passage, the cells were maintained in DMEM-F12 medium containing 4.5 g/l glucose, 10% FCS and penicillin/streptomycin.

### Histologic analysis - immunohistochemistry and immunofluorescence

Whole embryos and pancreas were fixed in 10% neutral buffered formalin and embedded in paraffin. For histopathological analysis, pancreas were serially sectioned (3 µm) and stained with hematoxylin and eosin (H&E). Histopathological analysis of pancreas and scoring of tissue damage, were performed using representative serial H&E-stained sections (50 µm apart, at least two sections per pancreas). Immunostainings followed standard protocols. Antibodies used for immunohistochemical studies are listed in Table 2. Slides were imaged using a Hamamatsu Nanozoomer 2 slide scanner (Hamamatsu Photonics) or a Panoramic 250 flash II Scanner. Images were analyzed with caseViewer or QPath software.

**Table 2.** Antibodies used in this study

| Antigen | Host | Dilution | References | Manufacturer |
| --- | --- | --- | --- | --- |
| TRIP12 | Rabbit | 1:1000 (IHC)<br>1:1000 (WB) | Cat# HPA036835, RRID :AB_2675340 | Sigma-Aldrich |
| TRIP12 | Rabbit | 1:1000 (IHC)<br>1:1000 (WB) | A301-814A RRID: AB_1264344 | Bethyl Laboratories |
| TRIP12 | Rabbit | 1:1000 (WB) | Bo Am Seo<sup>75</sup> | N/A |
| Cytokeratin 19<br>(Troma III) | Rat | 1:100(IHC) | Rolf Kemler (Max-Planck Institute,<br>Freiburg, Germany) | DSHB/NICHD/ The<br>University of Iowa |
| PTF1a | Rabbit | 1:1000 (IHC,IF) | Marlène Dufresne<sup>34</sup> | Eurogentec |
| E-Cadherin | Mouse | 1:500 (IHC, IF) | Cat# 610181, RRID:AB_397580 | BD Biosciences |
| PDX1 | Rabbit | 1:1000 (IHC, IF) | Cat# ab47267, RRID:AB_777179 | Abcam |
| Amylase | Rabbit | 1:1000 (IHC, IF) | Cat# A8273, RRID:AB_258380 | Sigma-Aldrich |
| Insulin | Rabbit | 1:20 (IHC, IF) | Cat#F/AN6735-5M | Biogenex |
| Glucagon | Mouse | 1:500 (IHC, IF) | Cat# G2654, RRID:AB_259852 | Sigma-Aldrich |
| Claudin 18 | Rabbit | 1:1000 (IF) | Cat#700178, AB-34H14L15 | Invitrogen |

WB, Western-blot; IF, immunofluorescence ; IHC, Immunohistochemistry.

Murine cell lines were seeded on cover slips in 6-well plates at a density of 5.10^5^ cells. After 48 hours, cells were fixed with paraformaldehyde 4% for 20 minutes, permeabilized with Triton 0.25% for 15 minutes and saturated with a PBS-BSA 5% solution for 1 hour. Cells were incubated with primary anti-Claudin 18 antibody (Table 2) diluted in a PBS-BSA 3% solution overnight at 4°C. After PBS washes, cells were incubated with AlexaFluor 555®-anti-rabbit secondary antibody (ThermoFisher) for two hours at room temperature. Nuclei were counterstained with DAPI (ThermoFisher). Cover slips were mounted on glass slides using DAKO fluorescent mounting medium (DAKO). Immunostaining was visualized using Z1 videomicroscope (Zeiss) and images were acquired using Zen software.

### RNA isolation - quantitative real-time PCR analysis

Total RNA was isolated from cell lines or acinar cells with Trizol® reagent (ThermoFisher) according to supplier’s instructions. RNA concentration and integrity were measured with a NanoDrop® ND-1000 spectrophotometer (ThermoScientific). The RevertAid® minus H (Thermofischer) kit was used according to the manufacturer’s recommendations with 2 µg of total RNA. qPCR duplicates were carried out using 2 µL of diluted cDNA (1 into 100) in a 20 µL-reaction using Sso Fast EvaGreen reagent (BioRad) following supplier’s instructions on a StepOne Plus™ Real-Time PCR system (Applied BioSystems). Relative amount of mRNA was calculated by the comparative threshold cycle (Ct) method as 2^-ΔCt^, where ΔCt=Ct target mRNA – Ct geometric mean of cyclophilin A and HPRT for cell lines or Ct geometric mean of HPRT, cyclophilin A et 18S for acini. Amplification efficacy was verified for every sets of primers, primers sequences are listed in Table 1.

### RNASeq analysis

Total RNA was isolated using Trizol® reagent (ThermoFisher) following manufacturer’s recommendations. Preliminary quality control of total mRNA was ensured by agarose gel electrophoresis and NanoDrop measurement. Sample quantitation and integrity were performed by Nanodrop and Agilent 2100, respectively. Paired-end (PE150) sequences (9 Go for cell lines and 15 Go for acinar cells) were generated using a NovaSeq 6000 apparatus (Illumina) by Novogen (Cambridge, UK).

Paired reads stored in fastq format were mapped on Mus Musculus reference genome (Mouse_GRCm39 version 2.7.4a) with STAR software (version 2.7.10a). Mapped counts for each individual mRNA in each sample were then processed on RStudio (version 4.2.0) with “DESeq2” library (version 1.36.0) for group comparison and “org.Mm.eg.db” library (version 3.16.0) for mRNA annotation. mRNA with either no available count in one sample, a count basemean lower than 100 and or an adjusted P-value from differential analysis higher than 0.05 were removed from subsequent analysis.

### Western-blotting

Murine cell lines were incubated in RIPA lysis buffer containing 50 mM Tris-HCl (pH 8.0), 150 mM NaCl, 0.1% SDS, 0.5% Sodium Deoxycholate, 1% Triton-X100® supplemented with Protease Inhibitor Cocktail (Sigma-Aldrich) for 20 minutes on ice, sonicated two times for 30 sec at 25% amplitude (Vibra-Cell sonicator) and centrifuged 20 minutes at 12 000 rpm Protein quantity was determined using Bradford assay (company à voir). Protein quantity was determined using Bradford assay (ThermoFischer). 30-50 µg of nuclear proteins were loaded onto a 10% SDS-PAGE and transferred on nitrocellulose membrane using TransBlot apparatus (BioRad). Membranes were subsequently incubated with TBS Tween-20® 0.01%, 5% fat milk for 1h and with indicated antibodies (listed in Table 2) overnight at 4°C. Membranes were incubated with appropriate HRP-coupled secondary antibodies. Protein abundance was visualized using Clarity™ Western ECL substrate, ChemiDoc™ XRS+ device and ImageLab™ software (BioRad).

### Cellular growth rate

KPC and KPCT tumors-derived murine cell lines were seeded at a density of 5.10^5^ cells/35mm-well. After 24h cells were counted every 24 hours for 72 hours using a Coulter a Beckman Z1 Coulter apparatus. Experiments were performed at least three times for each cell line.

### Colony formation assay

Cells were seeded at 10^3^ cells per 10 cm-plate. After 9-14 days, colonies were stained using 0.5% crystal violet. Images were acquired with videomicroscope cell observer (Zeiss) and analyzed using FiJi software. Experiments were performed three times for each cell line.

### Tail vein injection

Pancreatic cancer cells (1.10^6^) were injected into the tail vein of 8-weeks-old C57/B6 mice. Four-weeks post-injection, lungs were inflated then fixed and embedded in paraffin.

### Invasion assay

Upper compartment of a Boyden chamber (Falcon) was coated with growth factor reduced Matrigel matrix (Corning) and 3.10^5^ cells seeded in serum-free DMEM F-12 medium. Four hours after seeding, complete DMEM F-12 medium was added to the lower compartment to launch invasion. Twenty-four hours after, cells were fixed using 4% paraformaldehyde for 20 minutes and stained with crystal violet 0.5% for 20 minutes. Non-migrated cells were removed from the insert using a cotton swab. Images were acquired with videomicroscope cell observer (Zeiss) and analyzed using FiJi software. Experiments were performed three times for each cell line.

### Statistical analysis

Data were analyzed by 2-tailed, unpaired Student’s t-test or Kruskall-Wallis test using a multiple statistics Graph Pad Prism 9 software package. A difference was considered significant when p value was lower than 0.05. Gepia data were analyzed by Mantel-Cox test. Number of independent experiments is indicated in the figure legends. *, **, *** and **** indicate a p value <0.05, 0.01, 0.001 and 0.0001 respectively

All authors had access to the study data and had reviewed and approved the final manuscript.

## Abbreviations used in this paper

ADM: acinar-to-ductal metaplasia
AP-1: activating protein-1
ASXL1: additional sex combs-like1
bHLH: basic helix loop helix
CLDN18: claudin18
CREB: C-AMP response element-binding protein
DAPI: 4′,6-diamidino-2-phenylindole
EGF: epidermal growth factor
EMT: epithelial-to-mesenchymal transition
FBW7: F-box/WD repeat-containing protein 7
GEPIA: gene expression profiling interactive analysis
HECT: homologous to the E6-associated protein carboxyl terminus
HPRT: hypoxanthine phosphoribosyltransferase
IPMN: intraductal papillary mucinous neoplasm
KLF4: Kruppel-like factor 4
MAPK: mitogen-activated protein kinase
Nr5a2: nuclear receptor subfamily 5 group A member 2
PanIN: pancreatic intraepithelial neoplasia
PDAC: pancreatic ductal adenocarcinoma
PDX1: pancreatic and duodenal homeobox1
PKC: protein kinase C
PRC: polycomb repressive complexe
PTF1a: pancreas associated transcription factor 1a
RBPJ-L: recombining binding protein suppressor of hairless-like
RING: really interesting new gene
SOX: SRY-box transcription factor
SWI/SNF: switching defective/sucrose non-fermenting
TMA: tissue micro array
TRIP12: thyroid receptor interacting protein 12
UBR1: ubiquitin protein Ligase E3 component N-recognin 1.

